# Extracellular HBV RNAs are heterogeneous in length and circulate as virions and capsid-antibody-complexes in chronic hepatitis B patients

**DOI:** 10.1101/320994

**Authors:** Lu Bai, Xiaonan Zhang, Weixia Li, Min Wu, Jiangxia Liu, Maya Kozlowski, Liang Chen, Jiming Zhang, Yuxian Huang, Zhenghong Yuan

## Abstract

Extracellular HBV RNA has been detected in both HBV-replicating cell culture media and sera from chronic hepatitis B (CHB) patients, but its exact origin and composition remain controversial. Here, we demonstrated that extracellular HBV RNA species were of heterogeneous lengths, ranging from the length of pregenomic RNA to a few hundred nucleotides. In cell models, these RNAs were predominantly associated with naked capsids although virions also harbored a minority of them. Moreover, HBV RNAs in hepatitis B patients’ blood circulation were localized in unenveloped capsids in the form of capsid-antibody-complexes (CACs) and in virions. Furthermore, we showed that extracellular HBV RNAs could serve as template for viral DNA synthesis. In conclusion, extracellular HBV RNAs mainly consist of pgRNA or the pgRNA species degraded by the RNase H domain of the polymerase in the process of viral DNA synthesis and circulate as CACs and virions. Their presence in the blood circulation of CHB patients may be exploited to develop novel biomarkers for HBV persistence.

## Importance

Although increasing evidence suggests the presence of extracellular HBV RNA species, their origin and molecular forms are still under debate. In addition to the infectious virions, HBV is known to secrete several species of incomplete viral particles, including hepatitis B surface antigen (HBsAg) particles, naked capsids and empty virions during its replication cycle. Here, we demonstrated that extracellular HBV RNAs were associated with naked capsids and virions in HepAD38 cells. Interestingly, we found that unenveloped capsids circulate in the blood of hepatitis B patients in the form of capsids-antibody-complexes (CACs) and, together with virions, serve as vehicles carrying these RNAs molecules. Moreover, extracellular HBV RNAs are heterogeneous in length and represent either pregenomic RNA (pgRNA) or products of incomplete reverse transcription during viral replication. These findings provide a conceptual basis for further application of extracellular RNA species as novel biomarkers for HBV persistence.

## Introduction

Hepatitis B virus (HBV) is still a major global health problem with estimated 257 million people worldwide that are chronically infected with HBV (1). HBV, together with duck hepatitis B virus (DHBV) and several other related animal viruses belong to the hepadnaviridae family (2). HBV virion comprises of an outer envelope and an inner icosahedral nucleocapsid (NC) assembled by 240 copies of core protein (HBc) and packaged with a 3.2 kb partially double-stranded circular DNA genome (3–8). In addition to the virions, a large amount of incomplete viral particles such as hepatitis B surface antigen (HBsAg) particles, empty virions and naked capsids can also be released from cells in the process of virus replication (9). Subviral HBsAg particles are spherical or rodlike and are present in vast excess over virions in sera of CHB patients (2). Empty virions share the same structure as virions but are devoid of nucleic acids (10–14). Naked capsids, which exit cells via a different route from virions (15–17), have the same structure as NCs, but are either empty or filled with viral RNA and immature viral DNA (7, 11, 18–20).

In NC, pgRNA undergoes reverse transcription into minus-strand DNA followed by plus-strand DNA synthesis (2, 21–25). Intracellular NCs can be packaged with viral nucleic acids at all levels of maturation, including pgRNA, nascent minus-strand DNA, minus-strand DNA-RNA hybrids and relaxed circular DNA (RC DNA) or double-stranded DNA (DSL DNA) (5, 7). Only the NCs with relatively mature viral DNA (RC or DSL DNA) are enveloped and secreted as virions. HBV replicating cells can release empty core particles assembled from HBc proteins and NCs that contain various species of replicative intermediate nucleic acids into the culture supernatant. However, while free naked capsids could be readily detected *in vitro* (7, 11, 18–20), they are hardly found in the blood of HBV-infected patients (17, 26, 27).

Although, extracellular HBV RNA was detected in both *in vitro* cell culture system and in clinical serum samples, its origin and composition remain controversial. It was proposed that extracellular HBV RNA represents pgRNA localized in virions (28). However, HBV spliced RNA and HBx RNA were also detected in culture supernatant of HBV stably replicating cells as well as in sera of CHB patients (29, 30). In addition, extracellular HBV RNA was also suggested to originate from damaged liver cells (31), naked capsids or exosomes (11, 30). Hence, these extracellular RNA molecules have never been conclusively characterized. Here, we demonstrate that extracellular HBV RNAs are composed of viral RNAs heterogeneous in length ranging from pgRNA (3.5 knt) to RNA fragments with merely several hundred nucleotides. These RNA molecules are actually pgRNA, 3’ receding pgRNA fragments that have not been completely reverse transcribed to DNA and pgRNA fragments hydrolyzed by RNase H domain of polymerase in the process of viral replication. More importantly, extracellular HBV RNAs are mainly localized in naked capsids and in virions released from HBV replicating cells *in vitro* and also in CACs and virions circulating in blood of hepatitis B patients.

## Results

### Extracellular HBV RNAs are heterogeneous in length and predominantly integral to naked capsids instead of virions in HepAD38 cell culture supernatant

To ascertain the origin of extracellular HBV RNA, we first examined viral particles prepared from culture medium of an *in vitro* HBV-stably transduced cell line. Human hepatoma HepAD38 cell line was used in this study as it sustains vigorous HBV replication under the control of a tetracycline repressible cytomegalovirus (CMV) promoter (32). Total viral particles were concentrated and centrifuged over a 10–60 % (w/w) sucrose gradient. Most of subviral HBsAg particles, virions and empty virions were detected between fractions ranging from 8 to 13 (Fig. 1A, upper and middle panels) and naked capsids, detected only by anti-HBcAg but not anti-HBsAg antibodies, were found in fractions 14 to 17 (Fig. 1A, middle and lower panels). The majority of viral nucleic acids were detected in fractions between 11 and 18 (upper panel of Fig. 1B), which coincided with the fractions containing virions (fractions 11–13), naked capsids (fractions 15–18) and the mixture of these particles (fraction 14). Consistent with previous observations, HBV virions are packed with mature viral DNA (RC or DSL DNA) while naked capsids contain both immature single-stranded DNA (SS DNA) and mature viral DNA (Fig. 1B, upper panel). Moreover, Northern blot results showed that most of the HBV RNA was detected in the naked capsids (Fig. 1B, lower panel, fractions 15–18), whereas only a very small amount was associated with virions (Fig. 1B, lower panel, fractions 11–13). Contrary to reports describing extracellular HBV RNA as distinct pgRNA or the spliced HBV RNA species (29), HBV RNA detected in naked capsids ranged from the length of pgRNA down to a few hundred nucleotides (shorter than the HBx mRNA [0.7 knt]) and RNA molecules within virions are much shorter than within naked capsids. We excluded the possibility of artifact generated by the SDS-Proteinase K extraction method as similar RNA blot pattern was obtained using a TRIzol reagent to extract both intracellular nucleocapsid-associated and extracellular HBV RNA (Fig. S1). Also, quantification of viral RNA extracted by either SDS-Proteinase K method or TRIzol reagent produced a very close copy number, except that the TRIzol reagent is known to preferentially extract RNA rather than DNA (Fig. S1).

**FIG 1.**
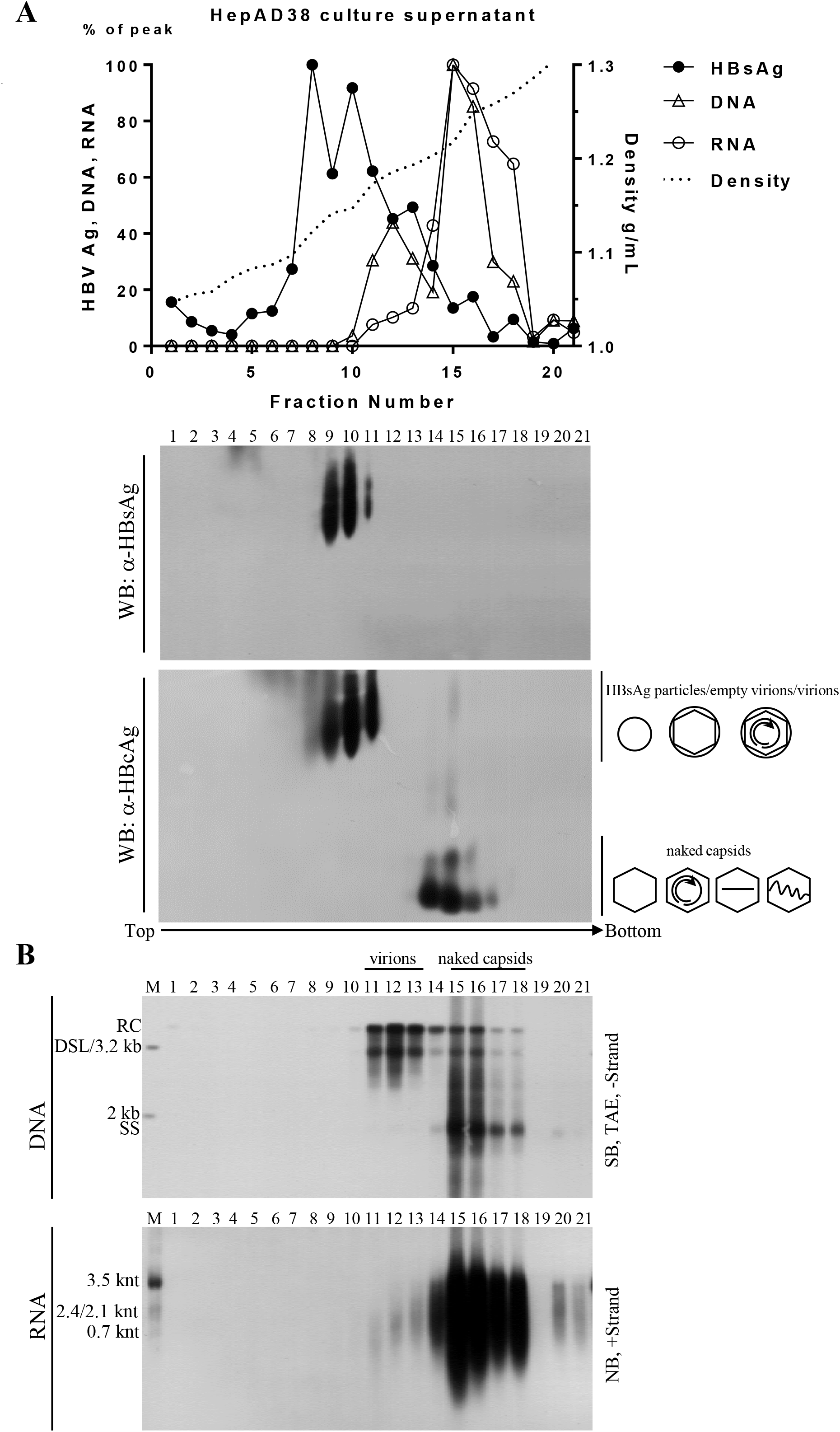
Sucrose gradient separation and analysis of viral particles from HepAD38 cell culture supernatant. (A) Distribution of hepatitis B viral particles-associated antigens and DNA/RNA in sucrose gradient. Viral particles prepared from HepAD38 cell culture supernatant (via PEG 8000 precipitation) were layered over a 10–60% (w/w) sucrose gradient for ultracentrifugation separation. Fractions were collected from top to bottom and HBsAg level was analyzed by enzyme-linked immunosorbent assay (ELISA). HBsAg and viral DNA and RNA (quantified from gray density of bands on (B)) signals and sucrose density were plotted together. Viral particles were first resolved by native agarose gel electrophoresis followed by Western blotting of HBV core and envelop proteins with anti-HBsAg and anti-HBcAg antibodies. (B) Detection of viral DNA/RNA by Southern or Northern blotting. Total viral nucleic acids were extracted by SDS-Proteinase K method and viral DNA and RNA (treated with DNase I) were detected by Southern and Northern blot analysis with minus- or plus-strand specific riboprobes, respectively. Symbols of HBsAg particles, empty virions (without nucleic acid), virions (with RC DNA) and naked capsids (empty or with nucleic acids) were depicted on the right side of lower panel of (A). Blank, no nucleic acids; two centered and gapped circles, RC DNA; straight line, SS DNA; wavy lines, pgRNA; M, markers (50 pg of 1 kb, 2 kb and 3.2 kb DNA fragments released from plasmids as DNA ladder or total RNA extracted from HepAD38 cells as RNA ladder).

To confirm the above results and to better separate naked capsids from HBV virions, isopynic CsCl gradient ultracentrifugation was employed. Naked capsids, detected only by anti-HBcAg, and not by anti-HBsAg antibodies, were mainly observed in fractions between 5 and 7 with densities ranging from 1.33 to 1.34 g/cm^3^ (Fig. 2A). The smearing bands of naked capsids were likely caused by high concentration of CsCl salt as fractionation of naked capsids in a 1.18 g/cm^3^ CsCl solution produced single bands on Western blots. Virions, detected by both anti-HBcAg and anti-HBsAg antibodies (Fig. 2A, upper and middle panels), were packaged with viral DNA (Fig. 2A, lower panel) and settled in fractions 13 to 15 with densities ranging from 1.23 to 1.25 g/cm^3^. In agreement with the results shown in Fig. 1, HBV virions contain only the mature viral DNA (RC or DSL DNA) while naked capsids contain viral DNA replicative intermediates that range from the nascent minus-strand DNA to mature viral DNA (Fig. 2B and C). The lengths of viral minus- and plus-strand DNA in naked capsids and virions were determined by alkaline agarose gel electrophoresis analysis as DNA fragments were denatured to single-strand molecules in alkaline condition and migrated according to their lengths. In contrast to the complete minus- and mostly complete plus-strand DNA (closed to 3.2 knt) in virions, both the minus-strand DNA and the plus-strand DNA can be complete and incomplete in naked capsids (shorter than 3.2 knt) (Fig. 2D and E). Moreover, the length of HBV RNAs within naked capsids still ranged from 3.5 knt of pgRNA to shorter than the 0.7 knt of HBx mRNA with pgRNA accounted for only 10% of total RNA signal detected by Northern blotting (quantified from gray density of bands in Fig. 2F). In contrast, virions are packed with relatively shorter and barely detectable level of HBV RNA species. Furthermore, naked capsids (fractions 3–7) and virions (fractions 10–21) were pooled and their respective viral DNA and RNA copy numbers were determined by quantitative PCR. Quantification results showed that viral DNA signals detected in naked capsids and in virions accounted for about 60% and 40%, respectively of total viral DNA signal in the HepAD38 cell culture supernatant (Fig. 2G). More importantly, 84% of the HBV RNA was associated with naked capsids, while merely 16% was detected within virions (Fig. 2G). Additionally, DNA/RNA ratio was eleven in virions and three in naked capsids (Fig. 2H), supporting the result that more HBV RNA is present in naked capsids.

**FIG 2.**
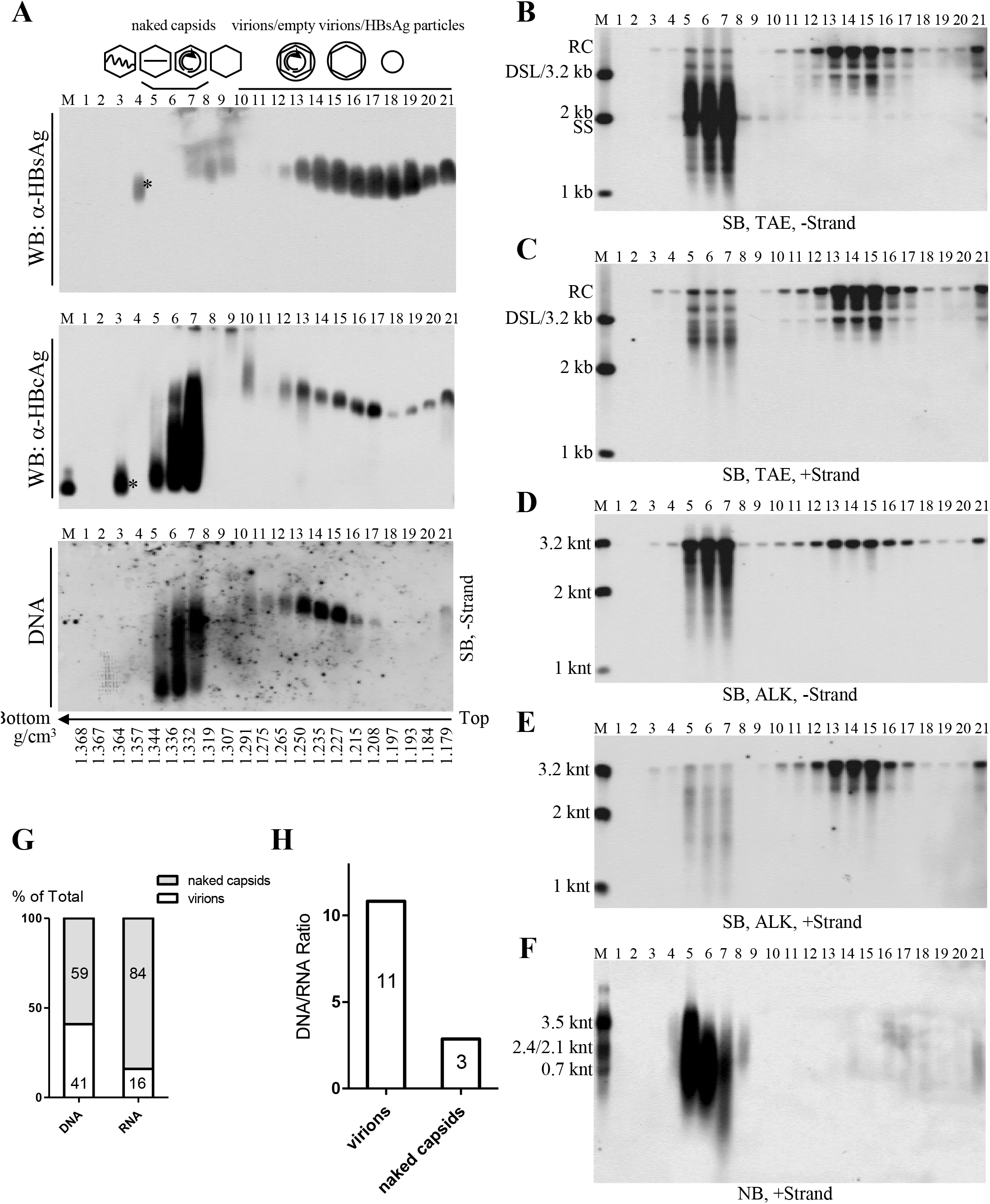
CsCl density gradient separation and analysis of viral particles from HepAD38 cell culture supernatant. (A) Native agarose gel analysis of viral particles. Culture supernatant of HepAD38 cells was concentrated (via ultrafiltration) and fractionated by CsCl density gradient centrifugation (3 ml of 1.18 g/cm^3^ CsCl solution in the upper layer and 1.9 ml of 1.33 g/cm^3^ CsCl solution in the lower layer). Viral particles in each fraction were resolved by native agarose gel electrophoresis followed by detection of viral antigens with anti-HBsAg and anti-HBcAg antibodies and viral DNA by Southern blotting with minus-strand specific riboprobe. (B-F) Southern and Northern blot detection of viral nucleic acids. Viral DNAs were separated by electrophoresis through Tris-Acetate-EDTA (TAE) or alkaline (ALK) agarose gel for Southern blotting with minus- or plus-strand specific riboprobes. Viral RNA was obtained by treating with total nucleic acids with DNase I and separated by formaldehyde/MOPS agarose gel followed by Northern blotting. (G) Quantification viral DNA and RNA in naked capsids or virions. Fractions contained naked capsids (fractions 3–7) or virions (fractions 10–21) were pooled and viral DNA and RNA were quantified by PCR method. (H) DNA and RNA ratio in naked capsids and virions calculated based on quantitative results. Asterisks (*) indicate unknown high density viral particles detected by anti-HBcAg or anti-HBsAg antibodies but devoid of any HBV-specific nucleic acids. M, markers (E. coli-derived HBV capsids or DNA and RNA ladders as described in Fig. 1).

### Extracellular HBV RNAs and immature viral DNA are detected in sera from CHB patients

Employing the HepAD38 cell culture system, we demonstrated the presence of extracellular HBV RNAs and immature and mature viral DNA packaged in both the naked capsids and virions. Interestingly, Southern blot analyses showed that SS DNA could also be observed in serum samples from some CHB patients. We speculated that SS DNA in circulation might be carried by capsid particles that were released by HBV-infected hepatocytes into patients’ bloodstream. However, we reasoned that due to strong immunogenicity of naked capsids (33, 34), it would be difficult to detect them as free particles but rather they would form complexes with specific anti-HBcAg antibodies and therefore circulate as antigen-antibody complexes (26, 33–35). To entertain this possibility, we then used protein A/G agarose beads to pull down the immune complexes. Our results showed that protein A/G agarose beads had greater specificity for the SS DNA-containing particles, circulating as presumed antibody-antigen complexes, than for the virions in patient’s sera (Fig. S2). As a result, forty-five serum samples obtained from CHB patients, with HBV DNA titer higher than 10^7^ IU per ml, were examined for the presence of particles containing SS DNA by a combination of protein A/G agarose beads pull-down assay and Southern blot analysis (Fig. 3A and B). Total HBV SS DNA was detected, albeit to different extents, in thirty-four serum samples (Fig. 3A and B, upper panels). The particles containing SS DNA were pulled down by protein A/G agarose beads from eleven out of the thirty-four samples (Fig. 3A and B, lower panels). Patients’ sera negative for SS DNA (patients 37, 38, 14 and 35) or positive for SS DNA (patients 17, 21, 42 and 44), as determined by the protein A/G agarose beads pull-down experiments, were selected for further studies.

**FIG 3.**
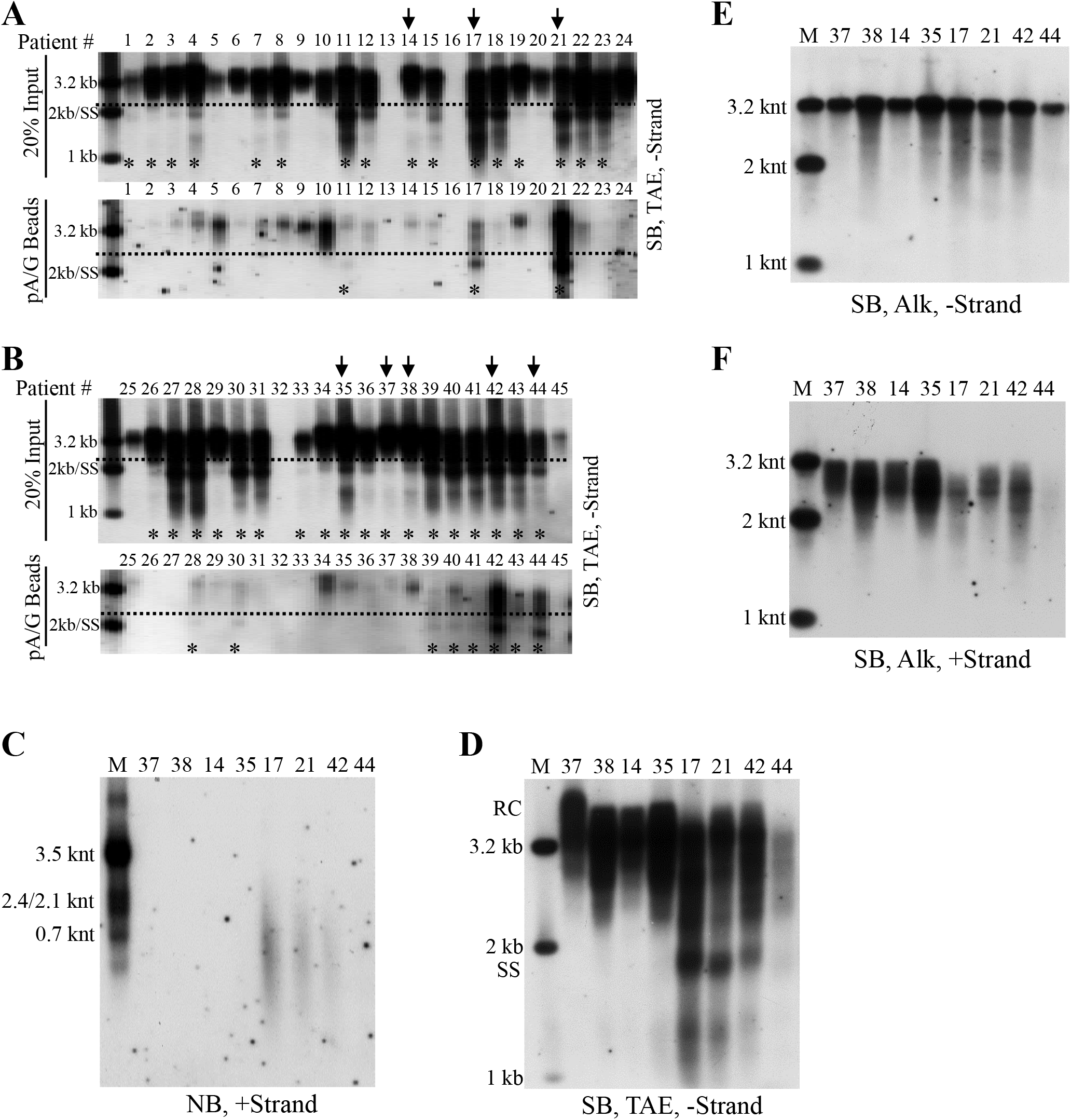
Characterization of HBV DNA and RNA in sera of CHB patients. (A-B) Analyses of serum viral DNA from CHB patients by Southern blotting. Viral DNA was extracted from serum samples obtained from forty-five chronic hepatitis B patients (20% of input sample used for protein A/G agarose beads pull-down) and subjected to Southern blot analysis. Alternatively, these samples were first incubated with protein A/G agarose beads and then viral DNA in the pull-down mixtures were analyzed by Southern blotting. Serum samples selected for further examining were marked with arrows and samples with SS DNA detection was labeled with asterisks (*). (C) Northern blot detection of serum viral RNA from patients 37, 38, 14, 35, 17, 21, 42 and 44. Total RNA were extracted from serum samples by TRIzol reagent and treated with DNase I before Northern blot analysis. (D-F) Southern blot analyses of viral DNA from selected samples. Viral DNA was separated by electrophoresis through TAE or alkaline agarose gels followed by Southern blot detection with indicated riboprobes.

Northern blot analyses showed that HBV RNA was only detected in serum samples from patients 17, 21 and 42 (Fig. 3C). Moreover, total viral DNA was analyzed by Southern blotting and SS DNA was readily observed in serum samples from patients 17, 21 and 42 (Fig. 3D). We also analyzed the lengths of viral minus- and plus- strand DNA in sera from these patients. Despite that most of minus-strand DNA was complete, a small amount of viral DNA (patients 38, 35, 17, 21 and 42) was shorter than 3.2 knt (Fig. 3E). Compared with viral minus-strand DNA, the length of plus-strand DNA, particularly in sera from patient 17, 21 and 42, was more variable and ranged from shorter than 2 knt to closed to 3.2 knt (Fig. 3F).

### Naked capsids form capsid-antibody-complexes (CACs) with anti-HBcAg antibody in blood circulation of CHB patients

We showed that SS DNA-containing particles were present in CHB patients’ sera. To further examine these particles, we used CsCl density gradient centrifugation to fractionate a serum mixture from patients 37, 38, 14 and 35. In agreement with our earlier results (Fig. 2A, lower panel, fractions 13–15) and previous reports, HBV virions, with the characteristic mature viral DNA (RC or DSL DNA), were detected in fractions from 12 to 14 with densities between 1.26 and 1.29 g/cm^3^ (Fig. 4A) (2). Careful inspection of the blots revealed that SS DNA, albeit at very low level, could be detected in fractions 8 and 9 with the densities from 1.33 to 1.34 g/cm^3^ and in fractions from 18 to 21 with the densities from 1.20 to 1.23 g/cm^3^ (Fig. 4A). In contrast, CsCl density gradient separation of viral particles from serum of the patient 17 showed a mixture of mature and immature viral DNA species. Thus, no distinct viral DNA (only mature RC or DSL DNA) specific to virions could be identified at densities between 1.27 and 1.29 g/cm^3^ while SS DNA was detected at broad densities ranging from 1.37 to 1.20 g/cm^3^ (Fig. 4B). Similar results were obtained using CsCl density gradient fractionation of sera from patient 21 and patient 46 (Fig. S3).

**FIG 4.**
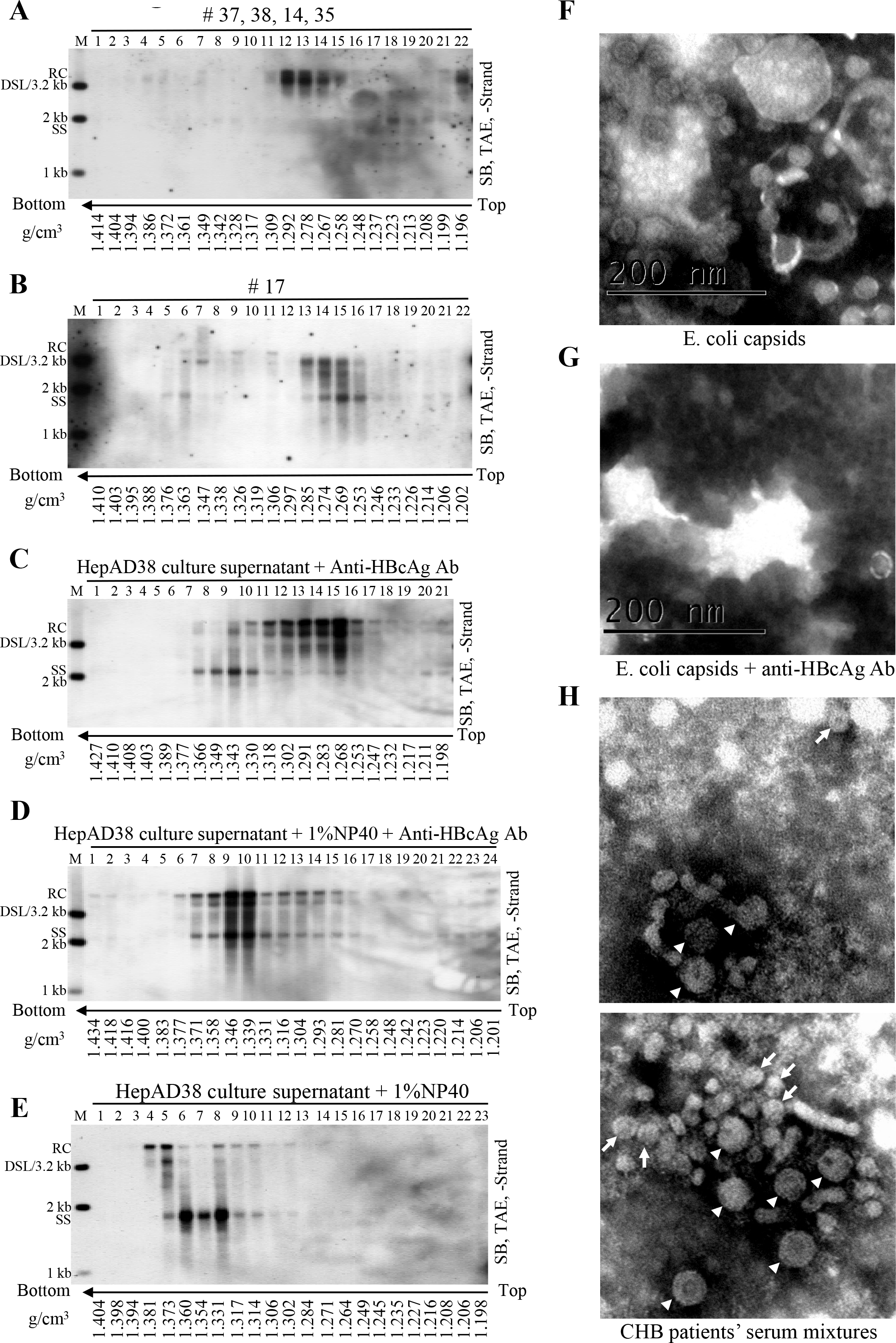
CsCl density gradient and EM analysis of hepatitis B viral particles. (A-B) CsCl density gradient analysis of viral particles in patient’s sera. One hundred microliter serum mixture from patients 37, 38, 14 and 35 (25 μL each) and 100 μL serum from patient 17 were separated by CsCl density gradient centrifugation (2 ml of 1.18 g/cm^3^ CsCl solution in the upper layer and 2.9 ml of 1.33 g/cm^3^ CsCl solution in the lower layer). Viral DNA in each fraction was extracted and detected by Southern blotting. (C-E) CsCl density gradient analysis of viral particles treated with detergent or anti-HBcAg antibody. Concentrated HepAD38 cell culture supernatant (250 μl each) (via ultrafiltration) was either mixed with anti-HBcAg antibody (10 μl) followed by incubated without or with NP-40 (final concentration: 1%) for 1 h at room temperature and 4 h on ice or treated with NP-40 only and then fractionated by CsCl density gradient ultracentrifugation. Viral DNA in each fraction was extracted and analyzed by Southern blotting. (F-G) EM of E. coli derived HBV capsids incubated without or with anti-HBcAg antibody. (H) EM of viral particles prepared from sera of CHB patients. Sera mixture (obtained from patients 11, 22, 23, 27, 28, 30 and 41) depleted of HBsAg particles were negatively stained and examined with electron microscope. The 42-nm HBV virions (arrowhead) and 27-nm naked capsids (arrow) were indicated while the smaller 22-nm rods and spheres of HBsAg particles could also be observed but not pointed out. Scale bars indicate 200 nm.

We proposed that naked capsids could be released into blood circulation of CHB patients but were neutralized by specific antibodies. As SS DNA was detected in both high and lower density region in CsCl gradient (Fig. 4B), we envisaged that the binding with specific antibodies led to a change of capsids’ buoyant density. To test this, anti-HBcAg antibody were mixed with HepAD38 cell culture supernatant to mimic the postulated CACs in serum samples. The results demonstrated that in contrast to SS DNA from naked capsids, distributed to three fractions at densities between 1.33 and 1.34 g/cm^3^ (Fig. 2A, lower panel, 2B), the mixture of naked capsids and CACs (SS DNA) distributed more widely and could be detected in the lower density region (1.25–1.32 g/cm^3^) (Fig. 4C, fractions 11–16).

To further confirm the lighter density of CACs, NCs in virions secreted to HepAD38 cell culture supernatant were treated with NP-40 and mixed with anti-HBcAg antibody. CsCl fractionation showed that naked capsids and virions-derived NCs have become a homogenous mixture banding at the densities from 1.37 to 1.27 g/cm^3^ (Fig. 4D). Likewise, virion-derived NCs obtained by treatment of serum sample from patient 46 with NP-40 together with original capsid-antibody complexes formed new homogeneous CACs that banded at densities between 1.23 and 1.27 g/cm^3^ (Fig. S3). However, NP-40 treatment alone did not produce a homogeneous mixture of naked capsids and virion-derived NCs as these two particles still settled at distinct density regions with their characteristic viral DNA (Fig. 4E). On the other hand, DNA molecules in the two types of capsids banded at densities between 1.38 and 1.31 g/cm^3^, further confirming that CACs have relatively lighter density.

Alternatively, the appearance of a homogenous mixture of virion-derived NCs and naked capsids (Fig. 4D) may suggest the formation of higher order antibody-mediated complexes of capsids. For instance, the complexes might not represent individual antibody-coated capsid particles but rather big CACs consisting of several capsid particles interconnected by antibodies. To verify whether intercapsid immune complexes exist, anti-HBcAg antibody was added into the purified HBV capsids expressed by E. coli and this mixture was examined by an electron microscope. E.coli-derived capsids were scattered as separated, distinct particles (Fig. 4F and Fig. S4A). However, addition of antibody caused capsids aggregation into clusters, making them too thick to be properly stained (Fig. 4G and Fig. S4B). In spite of this, few capsids, which may not be bound by antibodies or may be associated with antibodies but not forming the intercapsid antibody complexes could be observed by electron microscopy (EM) (Fig. 4G and Fig. S4B).

We then examined CACs in serum samples from CHB patients by EM. Sera from patients 11, 17, 21, 22, 23, 27, 28, 30 and 41 positive for SS DNA were combined. Serum mixture, depleted of HBsAg particles by centrifugation through a 20% and 45% (w/w) sucrose cushion, was examined by EM. The 27-nm capsid particles or CACs were visible (Fig. 4H, arrow) along with the 42-nm HBV virions (Fig. 4H, arrowheads) and the 22-nm spheres and rods of residual HBsAg particles (not indicated). However, the picture was not clear enough for us to conclusively determine if capsids were connected by or bound with antibodies as described for unrelated virus in *in vitro* experiments (36). In addition, it is possible that some of the CACs may not be visible by EM as the complexes maybe too thick to gain clear contrast between lightly stained and heavily stained areas (Fig. 4G and Fig. S4B).

Lastly, CACs might be heterogeneous, having different molecular sizes and isoelectric points (pI) in hepatitis B patients’ blood circulation. *In vitro* binding of HepAD38 cell culture supernatant-derived naked capsids with anti-HBcAg antibody changed their electrophoretic behavior and made them unable to enter the TAE-agarose gel (Fig. S5A). Moreover, viral particles from sera of patients 0, 37, 38, 14, 35, 17, 21, 42 and 44 could not enter agarose gels prepared in TAE buffer but in buffer with higher pH value (10 mM NaCHO_3_, 3 mM Na_2_CO_3_, pH 9.4) and they appeared as smearing bands on Southern blots (Fig. S5B and C). Hence, the irregular electrophoretic behavior of these viral particles may result from changes in molecular size and pI value of capsid particles (pI = 4.4) following their association with specific immunoglobulin G (or other types of antibodies) having different pI values (pI of human IgG may range from 6.5 to 9.5) (37–40).

### Circulating HBV RNAs are of heterogeneous lengths and mainly associated with CACs and virions in hepatitis B patient’s plasma

To characterize HBV RNAs circulating in CHB patients’ sera, a plasma sample from patient 0 was studied. Similar to results obtained for serum samples from patients 17, 21 and 46 (Fig. 4B and Fig. S3A and B), viral DNA in the plasma sample of patient 0 was detected in a broad density range in CsCl gradient and no bands specific to HBV virions or naked capsids could be identified, indicating the presence of a mixture of virions and CACs (Fig. 5A).

**FIG 5.**
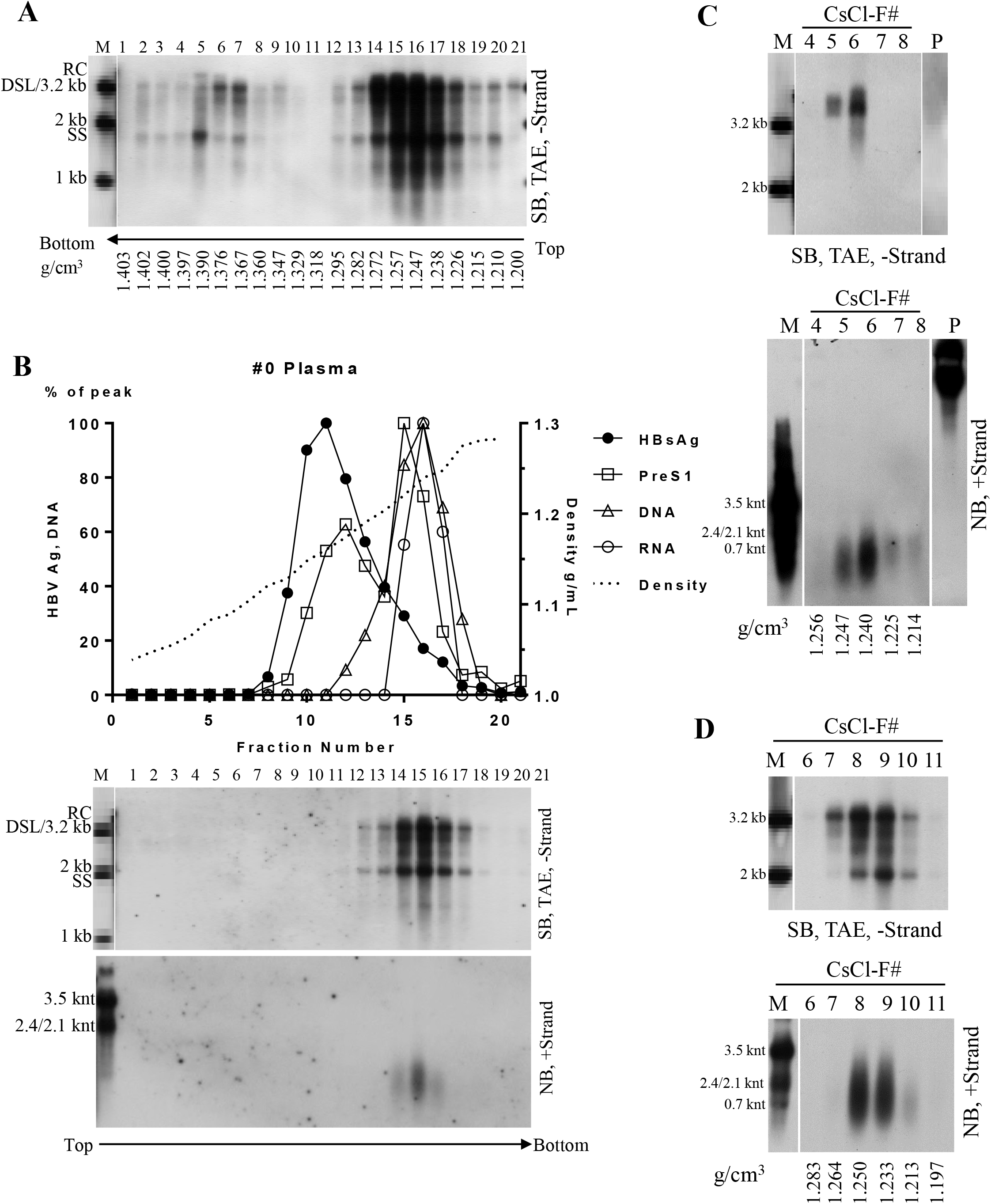
Characterization of nucleic acid content within viral particles in plasma sample from patient 0. (A) CsCl density gradient analysis of plasma sample. Plasma sample was not concentrated and added directly with CsCl salt to a concentration of 21% (w/w) or 34% (w/w). Two milliliter of the 21% CsCl-plasma mixture was underlayered with 2.9 ml 34% CsCl-plasma mixture followed by ultracentrifugation. Viral DNA from each fraction was extracted and subjected to Southern blot analysis. (B) Sucrose gradient analysis of concentrated plasma sample. Five hundred microliter concentrated plasma sample (via ultracentrifugation through a 20% sucrose cushion) was fractionated in a 10–60% (w/w) sucrose gradient. PreS1 and HBsAg levels were determined by ELISA. Viral DNA and RNA were detected by Southern and Northern blotting with minus- or plus-strand specific riboprobes. HBsAg, PreS1 and viral DNA and RNA (quantified from gray density of viral DNA/RNA bands in middle and lower panel of (B) signals and sucrose density were plotted together. (C) Analysis of concentrated plasma sample with lower CsCl density gradient centrifugation. 250μl of concentrated plasma sample was mixed with 2.2 ml TNE buffer and 2.45 ml of 37% (w/w) CsCl-TNE buffer (resulting a homogenous CsCl solution with density about 1.18 g/cm^3^) followed by ultracentrifugation. DNA in viral particle pellets (C, lane P) stuck to the sidewall of centrifugation tubes was recovered by digesting with SDS-Proteinase K solution. Viral DNA and RNA were subjected to Southern and Northern blot analysis. (D) Analysis of concentrated plasma sample with higher CsCl density gradient centrifugation. 250μl of concentrated plasma sample was mixed with one milliliter of TNE buffer and 1.25 ml of 37% (w/w) of CsCl-TNE buffer and underlayered with 2.4 ml of 27% (w/w) (1.25 g/cm^3^) CsCl-TNE solution followed by ultracentrifugation. HBV DNA and RNA was detected by Southern and Northern blotting.

Viral particles were first pelleted through a 20% sucrose cushion and then separated in a sucrose gradient. HBsAg was detected in fractions from 8 to 18, peaking at fraction 11. The preS1 antigen was found in fractions from 9 to 19 with the peak at fractions 12 and 15, indicating its presence in HBsAg particles and HBV virions (Fig. 5B, upper panel). Viral DNA, representing a combination of both mature and immature viral DNA, was detected in fractions from 13 to 18 (Fig. 5B, middle panel), suggesting the localization of CACs and virions in these fractions. HBV RNA was detected between fractions 15 and 17 and appeared in the same peak as viral DNA (Fig. 5B, lower panel), indicating that HBV RNA may be incorporated in the same viral particles as viral DNA. Therefore, circulating HBV RNA may be localized within CACs and/or virions.

To better characterize HBV RNA in CACs and virions, plasma sample from patient 0 was centrifuged through a 20% sucrose cushion and pellets were fractionated in a homogenous CsCl solution (1.18 g/cm^3^) as previous described (8). However, possibly due to a tendency of capsid particles to aggregate and stick to the wall of centrifugation tube and low density of the initial CsCl solution (8, 41), only mature DNA species from virions were detected in densities ranging from 1.22 to 1.24 g/cm^3^ (Fig. 5C, upper panel). Northern blot analyses demonstrated that the lengths of virion-associated HBV RNAs were about several hundred nucleotides (Fig. 5C, lower panel). Virion-associated RNAs were unlikely to be contaminated by CAC-associated HBV RNAs since the immature SS DNA could not be observed even after a long exposure and if CACs were present RNA should be longer (Fig. 5D, lower panel). Viral nucleic acids in pellets adhered to the centrifugation tube sidewall were retrieved and could be readily detected on Northern (Fig. 5C, lower panel, lane P) or Southern blots using plus-strand specific rather than minus-strand specific riboprobe (Fig. 5C, upper panel, lane P).

To analyze viral nucleic acids in CACs, concentrated plasma sample was separated in a higher CsCl density gradient (1.18 g/cm^3^ and 1.25 g/cm^3^). Both mature and immature viral DNA was only detected in fractions with densities from 1.21 to 1.26 g/cm^3^ (Fig. 5D, upper panel), indicating the presence of a mixture of HBV virions and CACs. Viral RNAs were detected and ranged from the length a little shorter than the full-length pgRNA to a few hundred nucleotides (Fig. 5D, lower panel). Comparing with virion-associated RNAs (Fig. 5C, lower panel), HBV RNA species detected in the mixture of CACs and virions were longer, with the longer RNA molecules possibly associated with CACs.

### Extracellular HBV RNAs could serve as templates for synthesis of viral DNA

Intracellular NCs are known to contain viral nucleic acids in all steps of DHBV DNA synthesis, including pgRNA, nascent minus-strand DNA, SS DNA and RC DNA or DSL DNA (5). Our results showed that naked capsids contain almost the same DNA replicative intermediates as that of intracellular NCs (Fig. 1B, 2B and F) (7, 11). We also demonstrated that extracellular HBV RNAs within the naked capsids, CACs and virions were heterogeneous in length (Fig. 1B, lower panel; Fig. 2F, 5C and D). In the presence of dNTPs, viral RNA could be degraded and reverse transcribed into minus-strand DNA by the endogenous polymerase *in vitro* (5, 42, 43). Also, incomplete plus-strand DNA with a gap about 600–2100 bases, could also be extended by endogenous polymerase (44, 45). Based on these results we wished to examine whether extracellular HBV RNAs could serve as RNA templates for viral DNA synthesis and be degraded by polymerase in the process. As shown in Fig. 6, EPA treatment led to viral minus- (Fig. 6A and C) and plus-strand (Fig. 6B and D) DNA extension and, more importantly, HBV RNA signal reduction (Fig. 6E, lanes 4 vs 6 and lanes 8 vs 10) in extracellular viral particles from either culture supernatant of HepAD38 cells or plasma sample from patient 0. The efficiency of EPA in extracellular viral particles from HepAD38 cell culture supernatant appeared to be low. This was likely caused by the detection of both extended and unextended DNA strands by the hybridization method we employed rather than a method that would detect newly extended radioactively labeled DNA.

**FIG 6.**
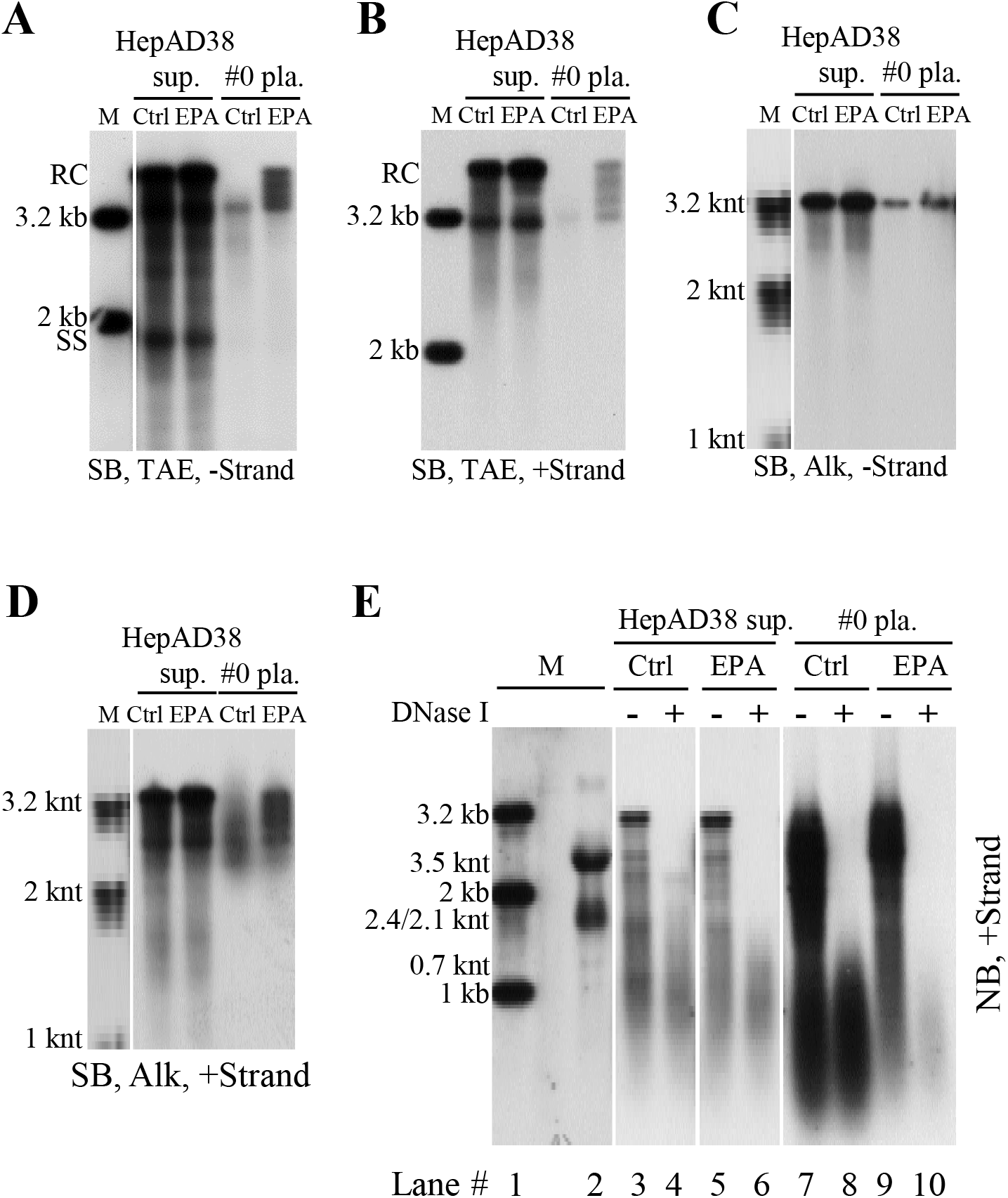
Analysis of extracellular HBV DNA and RNA by endogenous polymerase assay (EPA). (A-D) Southern blot analysis of viral DNA strand elongation after EPA treatment. EPA was carried out employing HepAD38 cell culture supernatant and plasma sample from patient 0. Total nucleic acids were extracted via SDS-Proteinase K method. Viral DNA was separated by electrophoresis in TAE or alkaline agarose gels followed by Southern blot analysis with minus- or plus-strand specific riboprobes. (E) Northern blot analysis of viral RNA changed upon EPA treatment. Total viral nucleic acids (lanes 3, 5, 7 and 9) or RNA (treated with DNase I) (lanes 4, 6, 8 and 10) were separated by formaldehyde/MOPS agarose gel electrophoresis and subjected to Northern blotting.

In the process of HBV DNA replication, prior to minus-strand DNA synthesis, capsid-associated RNA is the full-length pgRNA. Upon transfer of viral polymerase-DNA primer to 3’ DR1 region of pgRNA and cleavage of the 3’ epsilon loop RNA (a 3.2 knt-pgRNA fragment is remained), minus-strand DNA synthesis initiates and the pgRNA template is continuously cleaved from 3’ to 5’ by RNase H activity of viral polymerase. Consequently, from the initiation to the completion of minus-strand DNA synthesis, there will be a series of pgRNA fragments with receding 3’ ends ranging from 3.2 knt to 18 nt of the 5’ cap RNA primer (2, 22–25), representing the RNA templates that have yet not been reverse transcribed into minus-strand DNA. In addition, short pgRNA fragments resulting from cleavage by RNase H domain of polymerase will also be generated. Therefore, we used RNA probes spanning HBV genome to map whether these RNA molecules are present in extracellular naked capsids and virions.

Five probes that spanned HBV genome except for the overlapping region between 5’ end of pgRNA and RNA cleavage site (nt 1818–1930) were prepared to map the extracellular HBV RNAs from HepAD38 cell culture supernatant (Fig. 7A). Intracellular nucleocapsid-associated HBV RNA from HepAD38 cells was used as a reference. As the probes moved from 5’ end to 3’ end of pgRNA, especially for probes 1 to 4, RNA bands were shifting from wider range, including both short and long RNA species, to narrower range close to full-length pgRNA and fewer RNA species were detected (Fig. 7A, upper panel, lanes 2, 5, 8, 11, 14 and 17). Similarly, with the probes moving from 5’ end to 3’ end of pgRNA a stronger intensity band representing extracellular HBV RNAs detected by each probe, especially for probes 1 to 4, was also shifting toward longer RNA migration region (Fig. 7A, upper panel, lanes 3, 6, 9, 12, 15 and 18). It should be noted that the shifting pattern was more apparent when RNAs were detected with probes 1 to 4 but not probe 5. It is possible that the reverse transcription speed is relatively quicker in the initial reverse transcription step and, as a result, fewer pgRNA fragments will harbor RNA sequence within this region. Also, a short RNA species migrated faster than 0.7 knt in either intracellular nucelocapsids or naked capsids and virions could be detected by all probes (Fig. 7A, upper panel, lanes 2, 3, 5, 6, 8, 9, 11, 12, 14, 15, 17 and 18). These RNA molecules are likely represent the pgRNA fragments that have been hydrolyzed by RNase H domain of viral polymerase (including the 3’ epsilon loop RNA cleaved by polymerase in the reverse transcription step) (25). Collectively, as predicted, longer extracellular HBV RNA species that migrated slower and closer to the position of pgRNA had longer 3’ ends, whereas the shorter viral RNA molecules that migrated faster had relatively shorter 3’ ends, and the RNA species detected by all probes may possible represent products of pgRNA hydrolysis.

**FIG 7.**
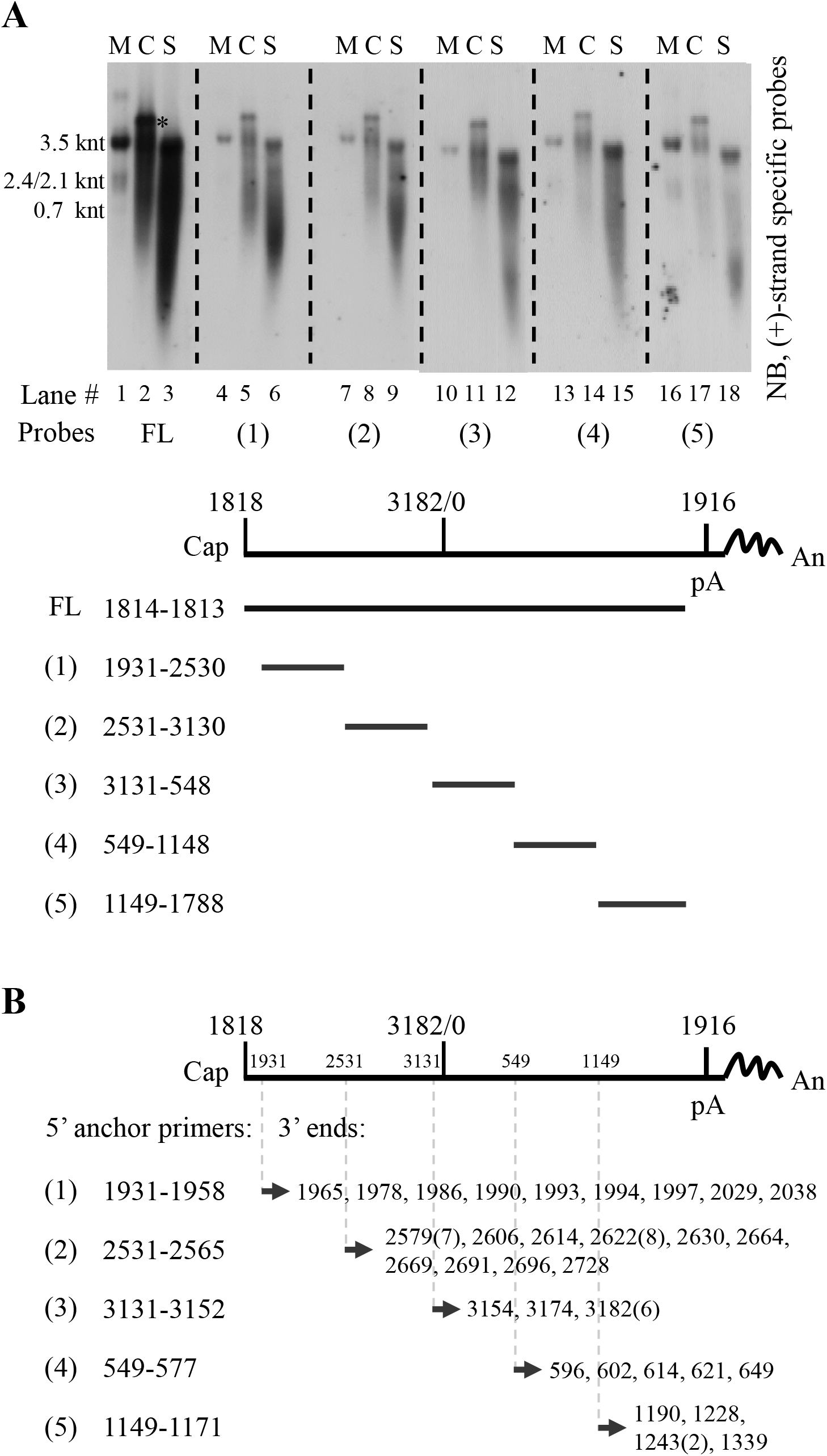
Mapping and identifying 3’ ends of extracellular HBV RNAs. (A) Northern blot detection of extracellular HBV RNAs with various riboprobes. Viral RNA from cytoplasmic (C) nucleocapsids (lanes 2, 5, 8, 11, 14 and 17) or culture supernatant (S) (lanes 3, 6, 9, 12, 15 and 18) of HepAD38 cells was extracted with TRIzol reagent and treated with DNase I before Northern blot analysis with plus-strand specific riboprobes spanning HBV genome as indicated. pgRNA was used as a reference and map coordinates were numbered according to sequence of HBV genome (genotype D, accession number: AJ344117.1). (B) Identification of 3’ ends of extracellular HBV RNAs. 3’ ends of extracellular HBV RNAs were identified by the 3’ RACE method using different HBV-specific anchor primers (the same 5’ primers used for generating templates for producing riboprobes used in [A, lower panel]). Identified 3’ ends were numbered as described above and numbers in brackets indicated amount of clones with the same 3’ ends. The asterisk (*) indicates unknown nucleic acid co-purified with intracellular capsid-associated viral RNA by TRIzol reagent. FL, full-length; Cap, 5’ cap of pregenomic RNA; pA, the polyadenylation site; An, poly(A) tail.

These results were further confirmed by employing a 3’ rapid-amplification of cDNA ends method. Various 3’ ends spanning HBV genome were identified (Fig. 7B), thus validating the presence of 3’ receding RNA and the heterogeneous nature of extracellular HBV RNAs.

EPA treatment clearly demonstrated that extracellular HBV RNAs could be used as templates for DNA synthesis and, the presence of 3’ receding end pgRNA fragments further confirmed not only the existence but also the use of such molecules as templates for viral DNA synthesis. Therefore, just like viral RNA counterpart within intracellular NCs, extracellular HBV RNA molecules represent the RNA molecules generated in the process of viral DNA replication.

### Entecavir reduces viral DNA level but increases extracellular HBV RNA level in naked capsids and virions *in vitro*

Entecavir (ETV), widely used in anti-HBV therapy, is a deoxyguanosine analog that blocks the reverse transcription and plus-strand DNA synthesis steps in HBV DNA replication process (46–48). Treatment of CHB patients with nucleos(t)ide analogs (NAs), including entecavir, efficiently reduces the level of serum viral DNA, but at the same time increases circulating HBV RNA level (29, 49–53). We examined the effect of entecavir on the levels of both intracellular and extracellular viral nucleic acids in HepAD38 cell culture.

Total viral RNA level remained unchanged or marginally increased upon entecavir treatment (Fig. 8A) and intracellular capsid-associated viral RNA level was increased (Fig. 8B, upper panel). In contrast and as expected, intracellular capsid-associated viral DNA level was decreased (Fig. 8B, lower panel). Similarly, extracellular viral DNA synthesis was significantly inhibited, while viral RNA was increased (Fig. 8C and D). Quantitative results showed that entecavir suppressed extracellular viral DNA to about one tenth but at the same time increased viral RNA about two folds of un-treated group (Fig. 8E).

**FIG 8.**
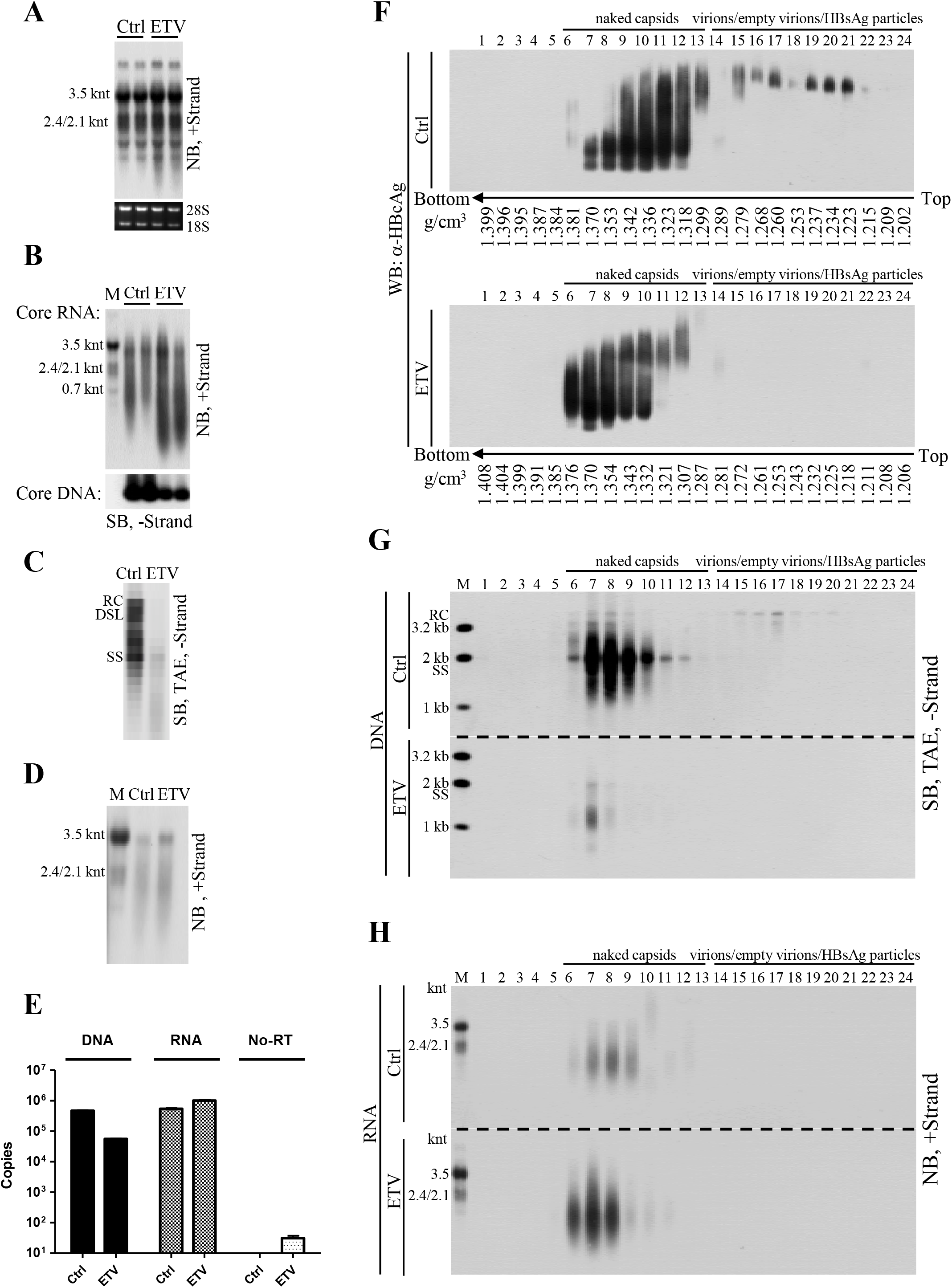
Analysis of HBV DNA and RNA change upon entecavir treatment of HepAD38 cells. (A) Change of total cellular HBV RNA level upon entecavir (ETV) treatment. HepAD38 cells were treated with entecavir (ETV) (0.1 μM) for 4 days and total cellular RNA was analyzed by Northern blotting with ribosomal RNAs serving as loading controls. (B) Change of intracellular nucleocapsid-associated viral RNA (core RNA) and DNA (core DNA) level after ETV treatment. Cytoplasmic core RNA was extracted by SDS-Proteinase K method and analyzed by Northern blotting. Intracellular nucleocapsids were first separated by native agarose gel electrophoresis and capsid-associated viral DNA (core DNA) was then probed with minus-strand specific riboprobe. (C-E) Change of extracellular HBV DNA and RNA level upon ETV treatment. Total nucleic acids in HepAD38 cell culture supernatant were extracted and subjected to Southern and Northern blot analysis with specific riboprobes or quantification by PCR method. (F-H) CsCl density gradient analysis of viral DNA/RNA level in naked capsids and virions after ETV treatment. HepAD38 cells were un-treated or treated with ETV and culture media were concentrated by ultrafiltration followed by fractionation in CsCl density gradients as described in Fig. 4. Viral particles in each fraction were separated by native agarose gel electrophoresis followed by Western blot analysis with anti-HBcAg antibody. Viral DNA and RNA were extracted and subjected to Southern or Northern blot analysis.

Since viral DNA and RNA were enclosed in both naked capsids and virions, CsCl density gradient was then used to separate these particles and to further study the antiviral effect of entecavir. As shown in Fig. 8, DNA-containing naked capsids were detected in fractions from 6 to 11 and virions in fractions between 15 and 24 (Fig. 8F). Entecavir effectively reduced viral DNA (Fig. 8G, fractions 6–10 and 15–17 and Fig. S6, fractions 6–10 and 15–17) but increased viral RNA content mainly in naked capsids (Fig. 8H, fractions 6–9). Moreover, the increase in RNA content within naked capsids led to an increased density of naked capsids (Fig. 8F, fractions 6 and 11 of lower panel vs fractions 6 and 11 in upper panel).

## Discussion

The RNA molecules in either intracellular NCs or extracellular virions were reported more than three decades ago (5, 42, 43), and naked capsids were shown to carry pgRNA *in vitro* (9, 11). Recently, it was suggested that the extracellular or circulating HBV RNA could serve as surrogate marker to evaluate the endpoint of hepatitis B treatment (28, 31, 49–54). With this in mind and to facilitate its application as a novel biomarker for viral persistence, we studied the origin and characteristics of extracellular HBV RNA.

In the present study, we extensively characterized extracelluar HBV RNAs and we demonstrated that extracellular HBV RNAs were mainly enclosed in naked capsids as opposed to complete virions in HepAD38 cells (Fig. 1B, 2F). These RNAs were of heterogeneous lengths ranging from full-length pgRNA (3.5 knt) to a few hundred nucleotides. Furthermore, circulating HBV RNAs, also heterogeneous in length, were detected in blood of hepatitis B patients (Fig. 3C, 5C and D). Interestingly, the detection of HBV RNAs coincided with the presence of immature HBV DNA (Fig. 3C and D). Isopycnic CsCl gradient ultracentrifugation of RNA positive serum samples exhibited a broad range distribution of immature HBV DNA, which contrasted with the results obtained in HepAD38 cells (Fig. 2B, 4B and Fig. S3A). For the first time, we provided convincing evidence that unenveloped capsids containing the full spectrum of HBV replication intermediates and RNA species heterogeneous in length, could be detected in blood circulation of chronic hepatitis B patients.

In view of our results and literature reports (2, 22–25), the presence of extracellular HBV RNAs could easily be interpreted in the context of the HBV DNA replication model (Fig. 9A). Since naked capsids contain viral DNA at all maturation levels, they will also carry HBV RNA molecules originating from pgRNA, including full-length pgRNA prior to minus-strand DNA synthesis, pgRNA with 3’ receding ends and the pgRNA hydrolysis fragments. On the other hand, virions that contain only mature form of viral DNA species would likely bear only the hydrolyzed short RNA fragments remaining in the nucleocapsid (44). Likewise, the HBV RNAs species found in CACs are longer than in virions in sera of hepatitis B patients (Fig. 5D, lower panel vs Fig. 5C, lower panel). In line with this reasoning, treatment of HepAD38 cells with entecavir, reduced viral DNA in naked capsids and virions (Fig. 8C, E and G; Fig. S6) but at the same time increased HBV RNA content within naked capsids (Fig. 8H). This may be a result of the stalled activity of viral RT with concomitant shutdown of RNA hydrolysis (47, 55).

**FIG 9.**
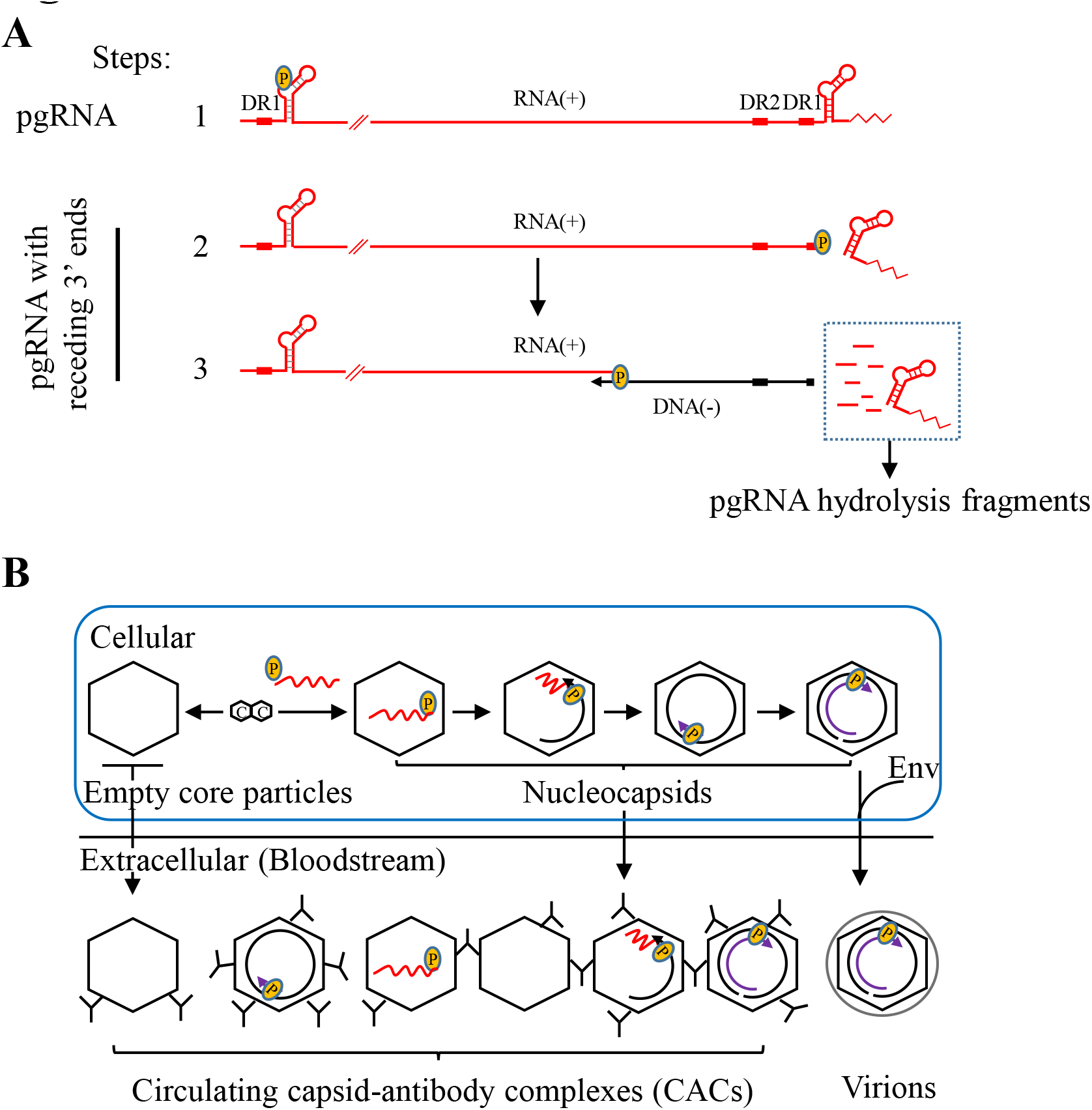
Models for the content of extracellular HBV RNAs and the formation of circulating CACs. (A) HBV RNA molecules present in the process of DNA synthesis. HBV RNAs are included in the following DNA synthesis steps: 1. Encapsidation of full-length pgRNA into NCs; 2. Transfer of polymerase-DNA primer to 3’ DR1 region and initiation of minus-strand DNA synthesis. 3’ epsilon loop of pgRNA will be cleaved by RNase H domain of polymerase; 3. Elongation of minus-strand DNA. With the extension of minus-strand DNA, pgRNA will be continuously cleaved from 3’ end generating pgRNA fragments with receding 3’ ends and pgRNA hydrolysis fragments. (B) Possible forms of circulating CACs. Empty core particles and intracellular NCs with pgRNA or pgRNA fragment and DNA replicative intermediates released into blood circulation of CHB patients are neutralized by specific antibodies (IgG), forming various forms of CACs.

Contrary to a recent report claiming that only the pgRNA-containing NCs can be enveloped and secreted as virions (28), we clearly demonstrated that secreted naked capsids carry the majority of HBV RNAs (Fig. 1B, 2F) and virion-associated RNAs are about several hundred nucleotides long (Fig. 1B and Fig. 5C). Our results are consistent with earlier reports demonstrating that only mature nucleocapsids with RC/DSL DNA are enveloped and secreted as virions (6–8, 11) and in this condition, virions carry only short RNase H-cleaved pgRNA (Fig. 9A, step 3).

In this research, we were unable to separate hydrolyzed pgRNA fragments from the pgRNA and pgRNA with 3’ receding ends. Thus, the length of these RNA molecules could not be determined. The existence of hydrolyzed RNA products during reverse transcription is not without precedent. In some retroviruses, DNA polymerization speed of RT is greater than RNA hydrolysis speed of RNase H, thus hydrolysis of RNA template is often incomplete (56, 57). For example, RT of avian myeloblastosis virus (AMV) hydrolyzed RNA template once for every 100 to 200 nucleotides (nts), while cleavage frequency of RTs of human immunodeficiency virus type 1 (HIV-1) and moloney murine leukemia virus (MoMLV) appeared to be around 100 to 120 nts (58). Moreover, RNA secondary structures, such as hairpins may stall the RT activity promoting RNase H cleavage producing shorter RNA fragments (56, 57).

Furthermore, the cleaved RNA fragments may not disassociate but anneal to the nascent minus-strand DNA forming the DNA-RNA hybrids until they are displaced by plus-strand DNA synthesis (56, 57). Although, similar studies on HBV replication were hampered by lack of functional viral polymerase *in vitro* (21, 59–61), the reported presence of DNA-RNA hybrid molecules clearly indicated the existence of degraded pgRNA fragments that still annealed to the minus-strand DNA (5, 42, 43, 62). Consistent with previous study, our results also showed that at least part of the SS DNA are associated with RNA molecules as the DNA-RNA hybrid molecules either by RNase H digestion or the cesium sulfate density gradient separation method as previously described (5).

Given the fact that HBV RNA and immature HBV DNA are packaged in naked capsids (Fig. 1B, 2B and F) (11), we postulated that, in CHB patients, unenveloped capsids are released into circulation where they rapidly form CACs with anti-HBcAg antibodies (Fig. 9B) (26, 34, 35). In support of this notion, we showed that: (1) Protein A/G agarose beads could specifically pull down particles with mature and immature HBV DNA from sera of CHB patients, implying the involvement of antibody (Fig. S2). (2) Addition of anti-HBcAg antibody to HepAD38 cell culture supernatant led to a shift of naked capsids’ buoyant density to lower density region (Fig. 4C and D), a pattern similar to that obtained in HBV RNA positive serum samples (Fig. 4B, 5A and Fig. S3A and B). (3) These particles exhibited higher pI and heterogeneous electrophoretic behavior, which differed from particles in HepAD38 culture supernatant, suggesting that they are not individual naked capsid particles but are associated with antibodies and take non-uniform compositions (Fig. 9B and Fig. S5) (37–39).

Apart from HBV particles, it was also reported that exosomes could serve as HBV DNA or RNA carriers (30, 63, 64). However, HBV DNA and RNA was exclusively detected in naked capsids or CACs and virions fractions rather than in lighter density regions where membrane vesicles like HBsAg particles (density of 1.18 g/cm^3^) and exosomes (density of 1.10–1.18 g/cm^3^) would likely settle (2, 28, 49, 65, 66). In addition, treatment of HepAD38 cell culture supernatant with micrococcal nuclease in the presence of detergent did not reduce the viral DNA and RNA signal, precluding the possibility that exosomes may serve as the main vehicles carrying HBV DNA or RNA molecules (Fig. S7).

In summary, we demonstrated that extracellular HBV RNA molecules are actually pgRNA and degraded pgRNA fragments generated in the HBV replication process. Moreover, we provided evidence that HBV RNAs exist in the form of capsid-antibody-complexes (CACs) in hepatitis B patients’ blood circulation. More importantly, the association of circulating HBV RNAs with CACs or virions in hepatitis B patients suggests their pgRNA origin. As pgRNA could only be transcribed from nuclear cccDNA (covalent closed circular DNA) instead of integrated HBV DNA fragments (67–69), its levels truly reflect the number or transcription status of cccDNA, especially for patients with lower serum HBV DNA titers when receiving NAs treatment. Hence, our results here strongly suggest the circulating HBV RNAs within CACs or virions in hepatitis B patients could serve as novel biomarkers to assess efficacy of treatment.

## Materials and methods

### Cell culture

HepAD38 cells that replicate HBV in a tetracycline repressible manner were maintained in Dulbecco’s Modified Eagle’s Medium (DMEM)-F12 medium supplemented with 10% fetal bovine serum and doxycycline was withdrew to allow virus replication (32).

### Patients and samples

Serum samples from forty-five chronic hepatitis B patients with HBV DNA titer higher than 10^7^ IU per ml were randomly selected. Detail medical records of these patients are included in Tab. S1.

Plasma sample was the plasma exchange product obtained from an HBeAg-negative hepatitis B patient (patient 0) (HBV genotype B with A1762T, G1764A and G1869A mutation) who died of fulminant hepatitis as a consequence of reactivation of hepatitis B (Tab. S1).

### Ethics statement

All the samples from HBV-infected patients used in this study were from an already-existing collection supported by National Science and Technology Major Project of China (Grant No. 2012ZX10002007–001). Written informed consent was received from participants prior to collection of clinical samples (70). Samples used in this study were anonymized before analysis. This study was conducted in compliance with the ethical guidelines of the 1975 Declaration of Helsinki and was approved by the ethics committee of the Shanghai Public Health Clinical Center.

### Preparation of viral particles

HepAD38 cell culture supernatant was added with polyethylene glycol (PEG 8000) to a final concentration of 10% (w/v) and incubated on ice for at least 1 h followed by centrifugation at 925g for 20 min. Pellets were suspended in TNE buffer (10 mM Tris-Cl [pH 7.5], 100 mM NaCl and 1 mM EDTA) containing 0.05% β-mercaptoethanol to 1/150 of original volume followed by brief sonication (71, 72). Alternatively, viral particles in HepAD38 cell culture supernatant were concentrated for 50–100 folds by ultrafiltration using a filter unit (Amicon Ultra-15, 100 KDa). Plasma sample from patient 0 were centrifuged through a 20% (w/v) sucrose cushion at 26,000 rpm for 16 h in an SW 32 Ti rotor (Beckman) and pellets were resuspended in 1/200 of original volume of TNE buffer and sonicated briefly (73).

Samples prepared using above methods were either used immediately or aliquoted and stored at –80°C for later use.

### Sucrose density gradient centrifugation

HepAD38 cells culture supernatant concentrated by PEG 8000 was centrifugation at 500g for 5 min to remove aggregates. 10%, 20%, 30%, 40%, 50% and 60% (w/w) sucrose gradients were prepared by underlayering and incubated for 4 h in waterbath at room temperature to allow gradient become continuous. Five hundred microliters of concentrated sample was layered over the gradient and centrifuged at 34,100 rpm for 14 h at 4°C in a Beckman SW 41 Ti rotor. Fractions were collected from the top to the bottom and density of each fraction was determined by refractometry (10). Fractions containing viral particles were subjected to native agarose gel analysis and HBsAg level was determined by enzyme-linked immunosorbent assay (ELISA) (Shanghai Kehua).

### Cesium chloride (CsCl) density gradient centrifugation

1.5 ml HepAD38 cells culture supernatant concentrated by ultrafiltration or serum samples from chronic hepatitis patients diluted with TNE buffer to 1.5 ml were mixed with equal volume of 37% (w/w) CsCl-TNE buffer (1.377 g/cm^3^) and underlayered with 1.9 ml 34% (w/w) CsCl-TNE buffer (1.336 g/cm^3^) followed by centrifuged at 90,000 rpm at 4°C for 12 h (Beckman VTi 90 rotor) (8). The tube was punctured from the bottom and every six to seven drops were collected as one fraction. Densities of separated fractions were determined by weighing. Each fraction was then desalted against TNE buffer by ultrafiltration followed by native agarose gel separation or nucleic acid extraction.

All the CsCl density gradient centrifugation experiments were carried out at 90,000 rpm at 4°C for 12 h in a Beckman VTi 90 rotor.

### Native agarose gel analysis of viral particles and capsid-associated DNA

Viral particles were resolved by native agarose gel (0.8% agarose gel prepared in Tris-Acetate-EDTA [TAE] buffer) electrophoresis and transferred in TNE buffer to either a nitrocellulose membrane (0.45 μM) for Western blot analysis of viral antigens or a nylon membrane for Southern blot analysis of viral DNA. For Western blotting, the membrane was first fixed as previously described (72) and HBV core antigen was detected by anti-HBcAg antibody (Dako) (1:5000); then, the same membrane was soaked in stripping buffer (200 mM Glycine, 0.1% SDS, 1% Tween-20, pH 2.2) and reprobed with anti-HBsAg antibody (Shanghai Kehua) (1:5000). For Southern blot analysis of viral DNA, the membrane was dipped in denaturing buffer (0.5 N NaOH, 1.5 M NaCl) for 10 seconds and immediately neutralized in 1 M Tris-Cl (pH 7.0)-1.5 M NaCl for 1 min followed by hybridization with minus-strand specific riboprobe (74).

### Viral nucleic acids extraction (I), separation (II) and detection (III)

**I**. To extract total viral nucleic acids (DNA and RNA), SDS-Proteinase K method was used (75). Samples were digested in solution containing 1% sodium dodecyl sulfate (SDS), 15 mM EDTA and 0.5 mg/ml proteinase K at 37°C for 15 min. Digestion mixture was extracted twice with phenol and once with chloroform. Aqueous supernatant were added with 1/9 volume of 3 M sodium acetate (pH 5.2) and 40 μg of glycogen and precipitated with 2.5 volumes of ethanol.

In addition to SDS-Proteinase K method, viral RNA was also extracted with TRIzol LS reagent according to manufacturer’s instructions (Thermo Fisher Scientific).

To isolate intracellular capsid-associated viral RNA, HepAD38 cells were lysed in NP-40 lysis buffer (50 mM Tris-Cl [pH7.8], 1 mM EDTA, 1% NP-40) and cytoplasmic lysates were incubated with CaCl_2_ (final concentration: 5 mM) and micrococcal nuclease (MNase) (Roche) (final concentration: 15 U/ml) at 37°C for 1 h to remove nucleic acids outside nucleocapsids. Reaction was terminated by addition of EDTA (final concentration: 15 mM) and then proteinase K (0.5 mg/ml without SDS) was added into the mixture followed by incubation at 37°C for 30 min to inactivate MNase (In contrast to DHBV capsids, HBV capsids are resistant to proteinase K digestion without SDS [personal observation]). Viral nucleic acids were released by addition of SDS to a final concentration of 1% and extracted as described above.

**II. (i) TAE agarose gel**. Viral DNA was resolved by electrophoresis through a 1.5% agarose gel in 1 x Tris-Acetate-EDTA (TAE) buffer followed by denaturation in 0.5 M NaOH-1.5 M NaCl for 30 min and neutralization with 1 M Tris-Cl (pH 7.0)-1.5 M NaCl for 30 min.

**(ii) Alkaline agarose gel**. Viral DNA was denatured with 0.1 volume of solution containing 0.5 M NaOH and 10 mM EDTA and resolved overnight at 1.5 V/cm in a 1.5% agarose gel with 50 mM NaOH and 1 mM EDTA. After electrophoresis, the gel was neutralized with 1 M Tris-Cl (pH 7.0)-1.5 M NaCl for 45 minutes (76).

**(iii) Formaldehyde/MOPS agarose gel**. Viral RNA was obtained by treatment of total nucleic acids extracted from above SDS-Proteinase K method with RNase free DNase I (Roche) for 15 min at 37°C. Reaction was stopped by addition of equal amount of 2 x RNA loading buffer (95% formamide, 0.025% SDS, 0.025% bromophenol blue, 0.025% xylene cyanol FF and 1 mM EDTA) supplemented with extra EDTA (20 mM) followed by denaturing at 65°C for 10 min. Viral RNA extracted by TRIzol LS reagent was mixed with 2×RNA loading buffer and denatured. Denatured mixtures were separated by electrophoresis through a 1.5% agarose gel containing 2% (v/v) formaldehyde solution (37%) and 1×MOPS (3-[N-Morpholino] propanesulfonic acid) buffer.

Above gels were balanced in 20×SSC solution (1×SSC is 0.15 M NaCl and 0.015 M sodium citrate, pH 7.0) for 20 min and viral nucleic acids were transferred onto nylon membranes overnight with 20×SSC buffer.

**III**. Digoxigenin-labeled riboprobes used for detection of HBV DNA and RNA were prepared by *in vitro* transcription of a pcDNA3 plasmid that harbors a 3215 bp of HBV DNA (nt 1814–1813) following vendor’s suggestions (Roche 12039672910). Riboprobes used for HBV RNA mapping were transcribed from DNA templates generated by PCR method by incorporating T7 promoter into the 5’ end of reversed primers (Tab. S2).

Hybridization was carried out at 50°C overnight followed by two five-minute washes in 2×SSC-0.1% SDS at room temperature and two additional fifteen-minute washes in 0.1×SSC-0.1% SDS at 50°C. The membrane was sequentially incubated with blocking buffer and anti-Digoxigenin-AP Fab fragment (Roche) at 20°C for 30 min. Subsequently, the membrane was washed twice with washing buffer (100 mM Maleic acid, 150 mM NaCl and 0.3% Tween-20, pH 7.5) for 15 min followed by detection with diluted CDP-Star substrate (ABI) and exposure to X-ray film.

### Protein A/G agarose beads pull-down of antibody-antigen complexes

Two hundred microliter of serum samples was first mixed with 300 μl of TNE buffer and then 15 μl of protein A/G agarose beads slurry (Santa Cruz) were added to the mixture followed by incubation overnight at 4°C in a sample mixer. Subsequently, protein A/G agarose beads were washed three time with TNE buffer and viral DNA in input serum samples (40 μl) and agarose beads pull-down mixtures were extracted and subjected to Southern blot analysis.

### Electron microscopy (EM)

Serum samples from patients 11, 17, 21, 22, 23, 27, 28, 30 and 41 were pooled (200 μl each) and mixed with 200 μl of 20% (w/w) sucrose. Serum mixtures were centrifuged through two milliliter of 20% (w/w) and two milliliter of 45% (w/w) (1.203 g/cm^3^) sucrose cushions at 34,100 rpm for 8 h at 4°C in an SW 41 Ti rotor (Beckman) to remove HBsAg particles. Supernatant were decanted and centrifugation tube was placed upside down for 20 seconds and residue sucrose was wiped out. One milliliter of phosphate buffer (10 mM Na_2_HPO_4_, 1.8 mM KH_2_PO_4_ and no NaCl) (pH 7.4) was added and the bottom of the tube was gently washed without disturbing the pellet. Then 11.5 ml of phosphate buffer was added into the tube and centrifuged again at 34,100 rpm for 3 h at 4°C. The pellet was resuspended in a drop of distilled water and dropped onto a carbon-coated copper grid followed by staining with 2% phosphotungstic acid (pH 6.1) and examining in an electron microscope (Philip CM120) (13, 77).

### Viral DNA and RNA quantification

Viral DNA used for quantification was extracted using SDS-Proteinase K method as described above. Viral RNAs were extracted by TRIzol LS reagent and DNase I was used to remove remaining DNA followed by phenol and chloroform extraction and ethanol precipitation. Reverse transcription were carried out using Maxima H Minus Reverse Transcriptase (Thermo Fisher Scientific) with specific primer (AGATCTTCKGCGACGCGG [nt 2428–2411]) according to manufacturer’s guidelines except the 65°C incubation step was skipped to avoid RNA degradation. Quantitative real time polymerase chain reaction (qPCR) was carried out using Thunderbird SYBR qPCR Mix (Toyobo) in a StepOnePlus real-time PCR System (ABI). Primer pairs (F: GGRGTGTGGATTCGCAC [nt 2267–2283]; R: AGATCTTCKGCGACGCGG [nt 2428–2411]) conserved among all HBV genotypes and close to 5’ end but not in the overlap region between the start codon and the polyA cleavage site of pgRNA were chosen. The cycling condition was 95°C for 5 min followed by 40 cycles of 95°C for 5s, 57°C for 20s and 72°C for 30s. DNA fragment containing 3215 bp of full-length HBV DNA was released from plasmid by restriction enzyme and DNA standards were prepared according to the formula that 1 pg of DNA equals to 3×10^5^ copies of viral DNA.

### Endogenous polymerase assay (EPA)

HepAD38 cells culture supernatant or plasma from patient 0 were concentrated as described above and mixed with equal volume of 2×EPA buffer (100 mM Tris-Cl pH 7.5, 80 mM NH_4_Cl, 40 mM MgCl_2_, 2% NP40 and 0.6% β-mercaptoethanol) with or without dNTPs (dATP, dCTP, dGTP and dTTP, each at a final concentration of 100 μM) (78). The reaction mixtures were incubated at 37°C for 2 h and stopped by addition of EDTA to a final concentration of 15 mM.

### 3’ rapid-amplification of cDNA ends (RACE)

Concentrated HepAD38 cell culture supernatant (by ultrafiltration) was digested with MNase in the presence of NP-40 (final concentration: 1%) for 30 minutes at 37°C. EDTA (final concentration: 15 mM) and proteinase K (final concentration: 0.5 mg/ml) were then added and incubated for another 30 min at 37°C. Viral nucleic acids were extracted with TRIzol LS reagent followed by DNase I treatment to remove residue viral DNA. Poly (A) tails were added to the 3’ end of HBV RNA by E. coli poly (A) polymerase (NEB). The pre-incubation step at 65°C for 5 min was omitted to reduce potential RNA degradation and reverse transcription were carried out with Maxima H Minus Reverse Transcriptase (Thermo Scientific) using an oligo-dT(29)-SfiI(A)-adaptor primer (5’-aagcagtggtatcaacgcagagt<u>ggccattacggcc</u>ttttttttttttttttttttttttttttt-3’) in reverse transcription buffer (1×RT buffer, RNase inhibitor, 1M Betanine, 0.5 mM each dNTP and 5 μM of oligo-dT(29)-SfiI(A)-adaptor primer) at 50°C for 90 min followed by heating at 85°C for 5 min and treatment with RNase H at 37°C for 15 min. PCR amplification of cDNA fragments were then performed with 5’ HBV-specific primers (the same sequences of forward primers used for riboprobes preparation (Tab. S2) except each primer containing a flanking sequence plus a SfiI(B) site [5’-agtgat<u>ggccgaggcggcc</u>-3’]) and 3’ adaptor primer (5’-aagcagtggtatcaacgcagagtg-3’). The reaction was carried out with PrimeSTAR HS DNA Polymerase (Takara) at 95°C for 5 min followed by 5 cycles of 98°C for 5s, 50°C for 10s and 72°C for 210s, 35 cycles of 98°C for 5s, 55°C for 10s and 72°C for 210s and a final extension step at 72°C for 10 min. PCR amplicons were digested with sfiI enzyme and cloned into pV1-Blasticidin vector (kind gift from Dr. Zhigang Yi in Shanghai Medical College of Fudan University). Positive clones were identified by sequencing and only clones with 3’ poly (dA) sequence were considered as the authentic viral RNA 3’ ends.

## Acknowledgements

We thank Zhuying Chen and Xiurong Peng for handling serum samples and compiling the clinical data used in this research.

---

**Suppl. Tab. 1:** Medical records of hepatitis B patients used in this research.

| Patient # | Sex | Age | HBV DNA<br>Titer<br>(IU/ml) | HBeAg<br>(IU/ml) | HBsAg<br>(IU/ml) | ALT<br>(IU/L) | SS<br>DNA |
| --- | --- | --- | --- | --- | --- | --- | --- |
| 0 | N/A | N/A | 2.67E+06 | - | 4932 | 396 | + |
| 1 | M | 54 | 1.24E+07 | 25 | >250 | 69 | + |
| 2 | F | 32 | 1.20E+07 | 1067 | 69384 | 38 | + |
| 3 | F | 21 | 1.36E+07 | 1712 | 200 | 149 | + |
| 4 | M | 33 | >5.00E+07 | 4812 | 113933 | 133 | + |
| 5 | N/A | N/A | 1.25E+07 | - | 3423 | 33 | - |
| 6 | M | 26 | 1.17E+07 | 545 | 2759 | 22 | - |
| 7 | M | 36 | 1.77E+07 | 4332 | 19541 | 136 | + |
| 8 | M | 35 | >5.00E+07 | 1199 | >250 | 104 | + |
| 9 | M | 26 | 2.20E+07 | - | >250 | 143 | - |
| 10 | M | 30 | >5.00E+07 | 2 | 4265 | 123 | - |
| 11 | F | 23 | >5.00E+07 | 20 | 5757 | 120 | + |
| 12 | M | 37 | 2.07E+07 | 2315 | 16128 | 177 | + |
| 13 | M | 28 | >5.00E+07 | 3495 | 60676 | 58 | N/A |
| 14 | F | 28 | >5.00E+07 | 16515 | 89575 | 78 | + |
| 15 | M | 37 | 1.62E+07 | 574 | +, ND | 112 | + |
| 16 | M | N/A | >5.00E+07 | 1601 | >250 | 22 | N/A |
| 17 | M | 15 | 2.28E+07 | 2038 | 32739 | 180 | + |
| 18 | M | 41 | 2.71E+07 | 694 | >250 | 313 | + |
| 19 | M | 34 | 2.35E+07 | 80 | 32514 | 148 | + |
| 20 | F | 44 | >5.00E+07 | 1596 | 4306 | 172 | - |
| 21 | M | N/A | 3.48E+07 | 107 | >250 | 103 | + |
| 22 | N/A | N/A | >5.00E+07 | 2024 | 45873 | 147 | + |
| 23 | M | 20 | 1.32E+07 | 13411 | 12387 | 344 | + |
| 24 | M | 48 | >5.00E+07 | 5511 | 76914 | 33 | - |
| 25 | M | N/A | 3.15E+07 | - | 15984 | 366 | - |
| 26 | M | 31 | 4.16E+07 | 10251 | 50469 | 442 | + |
| 27 | M | 60 | 1.35E+07 | 749 | >250 | 105 | + |
| 28 | F | 41 | >5.00E+07 | 4173 | >52000 | 194 | + |
| 29 | N/A | N/A | >5.00E+07 | 4233 | 49125 | 39 | + |
| 30 | M | 29 | 1.42E+07 | 25 | 5800 | 940 | + |
| 31 | M | 27 | 2.34E+07 | 1117 | 22412 | 129 | + |
| 32 | M | 37 | 2.65E+07 | - | 70 | 109 | N/A |
| 33 | N/A | N/A | 2.03E+07 | - | 4902 | 111 | + |
| 34 | M | 32 | >5.00E+07 | 993 | 43582 | 249 | + |
| 35 | N/A | N/A | 2.94E+07 | 4641 | 93336 | 12 | + |
| 36 | N/A | N/A | >5.00E+07 | 10956 | 2496 | 108 | + |
| 37 | F | 43 | >5.00E+07 | 1021 | >250 | 74 | + |
| 38 | F | 28 | >5.00E+07 | 215 | 446 | 26 | + |
| 39 | M | 31 | >5.00E+07 | +, ND | 38165 | 194 | + |
| 40 | N/A | N/A | >5.00E+07 | 25 | >250 | 69 | + |
| 41 | M | 26 | 1.52E+07 | +, ND | +, ND | 95 | + |
| 42 | M | 25 | >5.00E+07 | 6300 | 43151 | 373 | + |
| 43 | M | 22 | >5.00E+07 | 3844 | 23620 | 329 | + |
| 44 | M | 27 | 1.36E+07 | 1185 | 11106 | 149 | + |
| 45 | M | 44 | 1.28E+07 | 663 | 23330 | 425 | - |
| 46 | F | 29 | >5.00E+07 | +, ND | +, ND | 667 | + |

N/A: not available; ND: not determined; +: positive; -: negative; sera from patients 0 and 46 were not included with sera from other patients for SS DNA screening.

**Suppl. Tab. 2:**
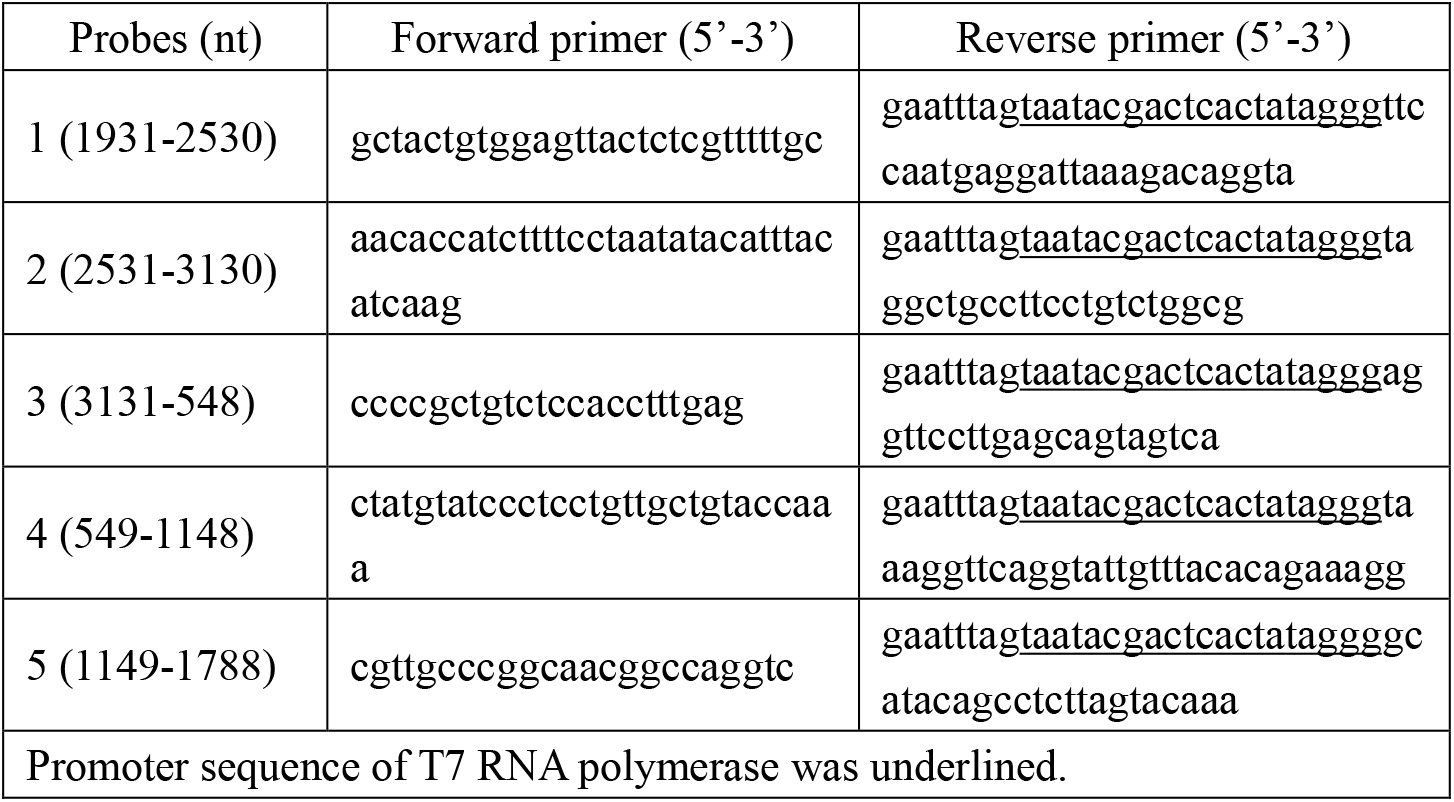
Primers used for preparing DNA templates for riboprobes transcription in extracellular HBV RNA mapping experiment.

**Suppl. Fig. 1:**
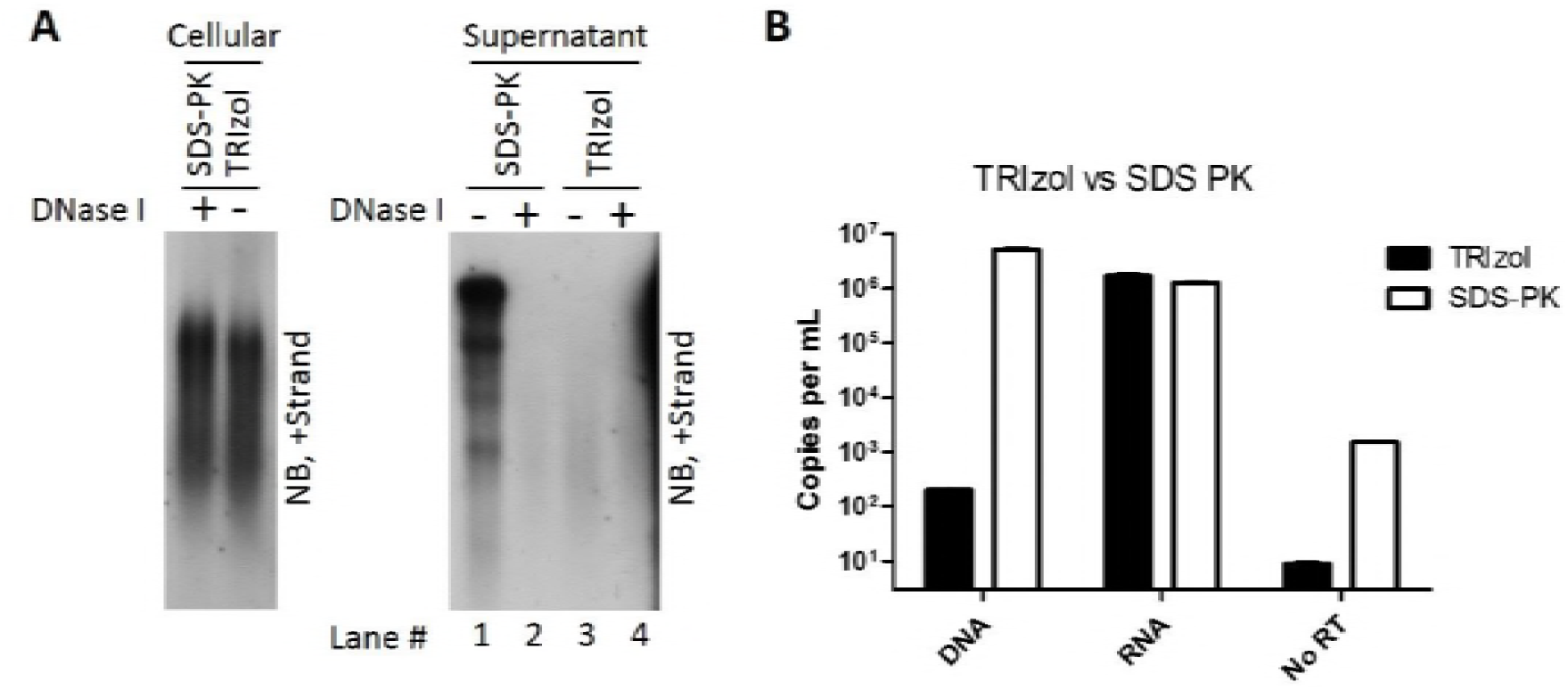
Comparison of viral DNA and RNA extraction efficiency of SDS-proteinase K and TRIzol reagent methods. Intracellular nucleocapsids were prepared from HepAD38 cells. Capsid-associated viral RNA was extracted by SDS-PK method (treated with DNase I [final concentration: 2 U/μl] to obtain RNA) or TRIzol reagent followed by Northern blotting (A, left panel). Total viral nucleic acids (A, right panel, lanes 1 and 3) or viral RNA (A, right panel, lanes 2 and 4) (treated with DNase I to remove DNA) prepared from culture supernatant of HepAD38 cells by above two methods were subjected to Northern blot analysis; meanwhile, viral DNA and RNA were quantified by PCR method (B).

**Suppl. Fig. 2:**
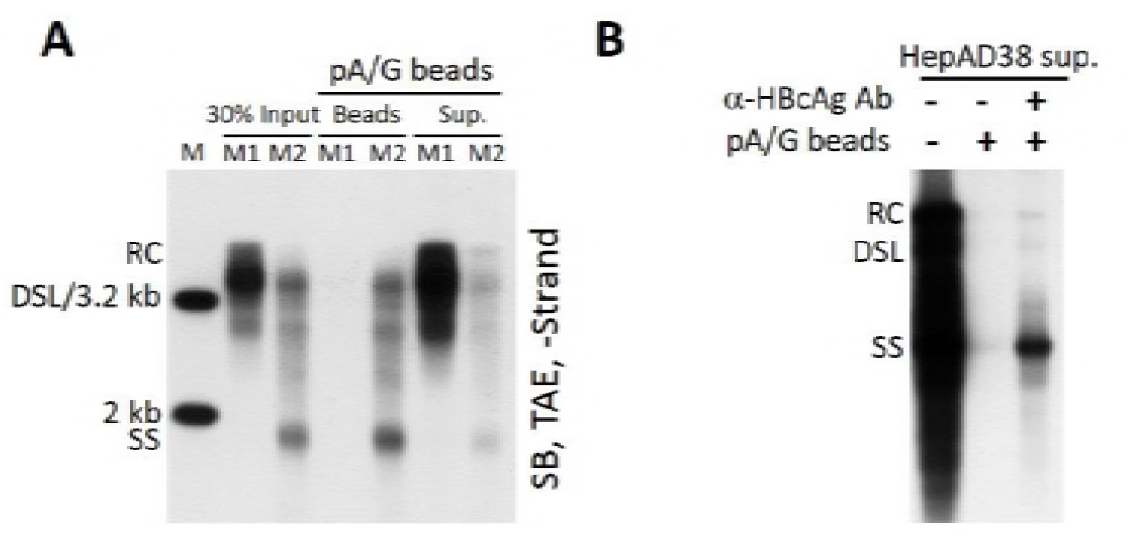
Analysis of the binding specificity of protein A/G agarose beads to capsid-antibody complexes (CACs) and to HBV virions. Sera (25 μl each) from CHB patients 37, 38, 14 and 35 (M1: mixture one) or from patients 17, 21, 42 and 44 (M2: mixture two) were pooled and incubated with 20 μl of protein A/G agarose bead slurry overnight at 4°C in a sample mixer. Viral DNA in input sera, protein A/G beads pull-down mixtures (beads) and the remaining supernatants (sup.) was extracted and subjected to Southern blot analysis (A). Concentrated HepAD38 cell culture supernatant (40 μl) was mixed with one milliliter of fetal bovine serum (FBS) and 20 μl of protein A/G agarose bead slurry with or without the presence of anti-HBcAg antibody. Viral DNA was extracted from input sample and protein A/G agarose beads pull-down mixtures followed by Southern blotting (B).

**Suppl. Fig. 3:**
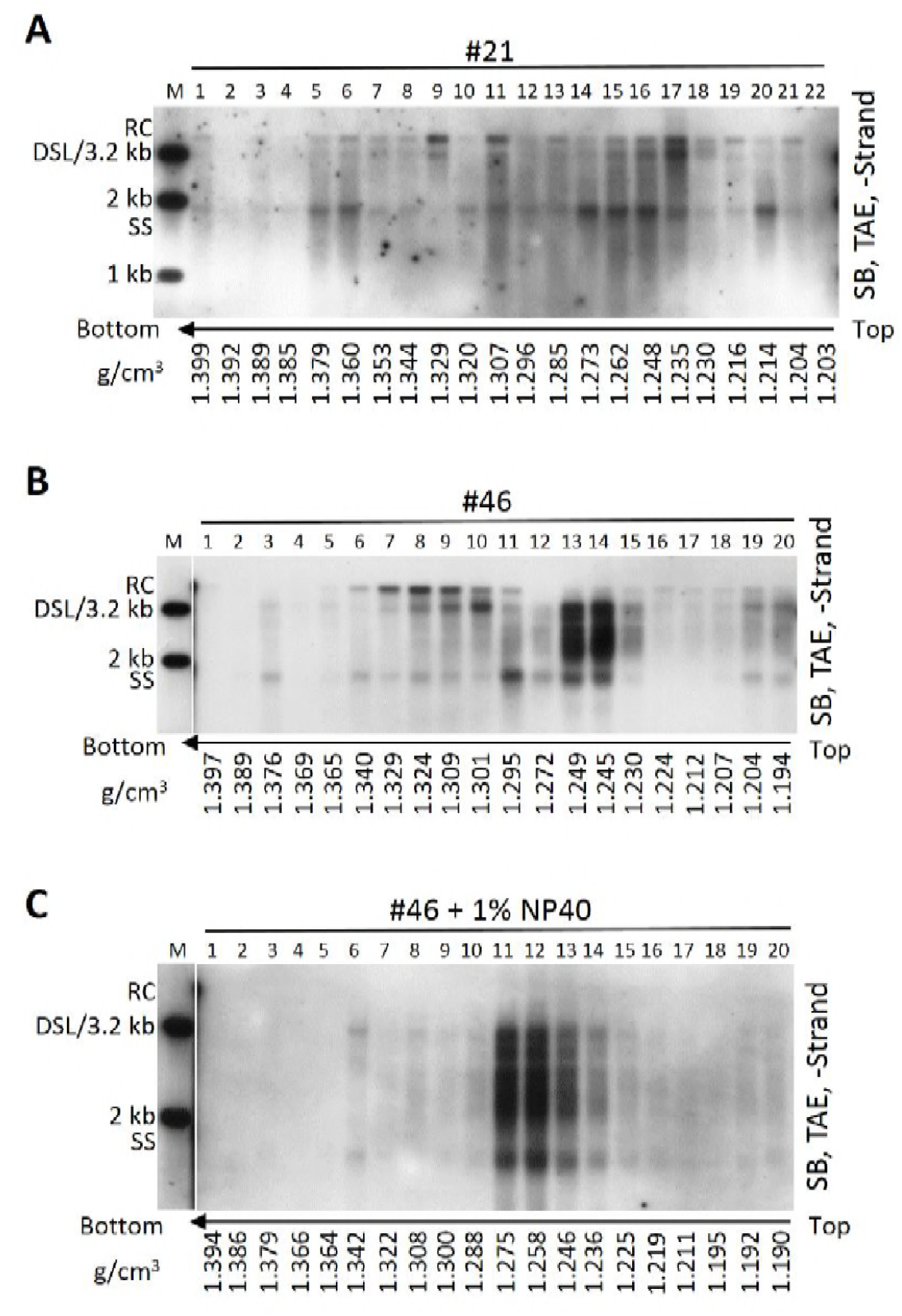
CsCl density gradient analysis of viral particles in sera of CHB patients. Serum from CHB patient 21 (A) or serum sample from patient 46 untreated (B) or treated (C) with NP-40 (final concentration: 1%) were fractionated by CsCl density gradient ultracentrifugation as described in Fig 4. Viral DNA in each fraction was extracted and subjected to Southern blot analysis.

**Suppl. Fig. 4:**
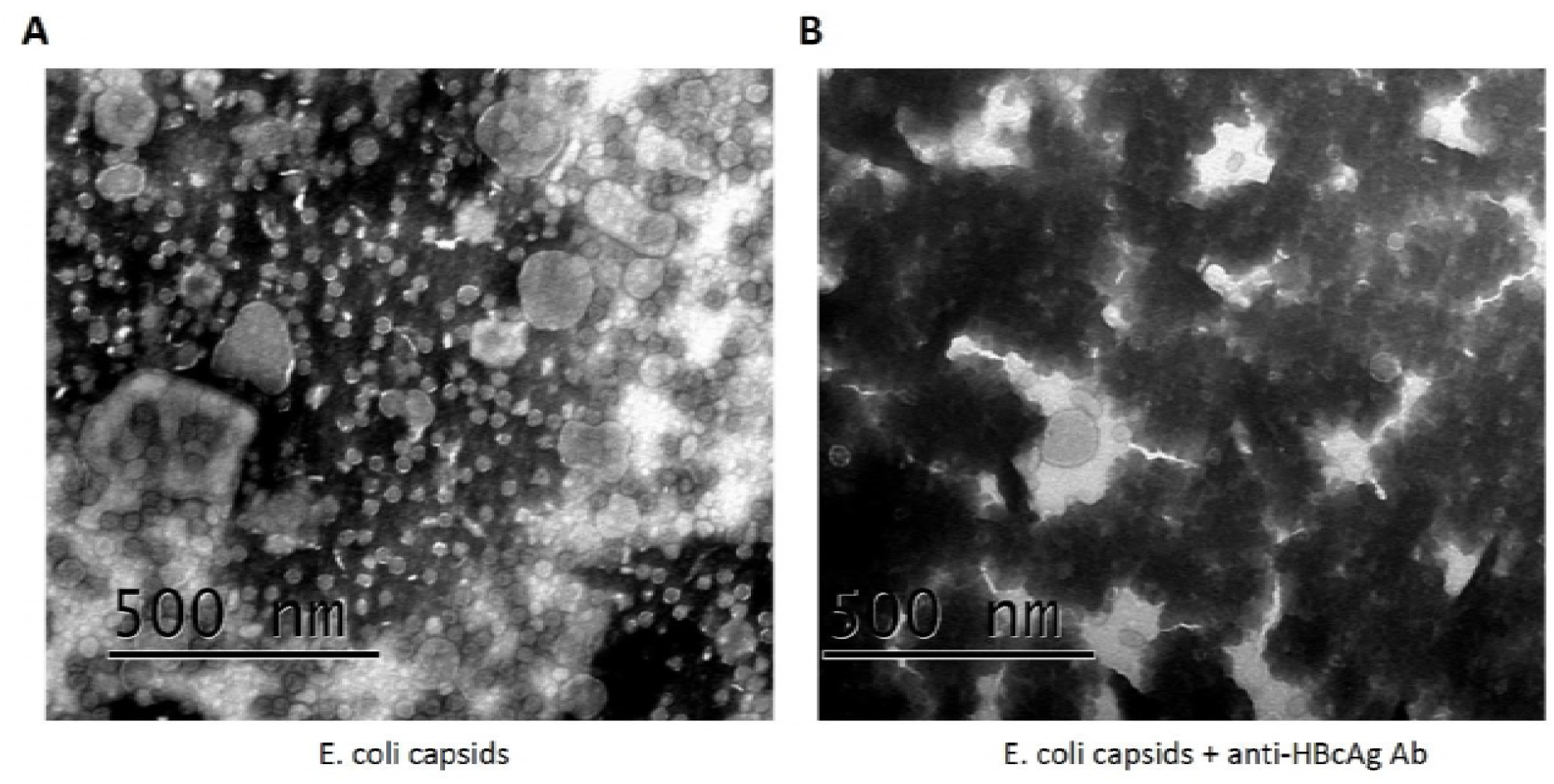
EM analysis of CACs formed by E. coli-derived capsids. HBV capsids were expressed and purified from E. coli. The capsids were incubated without (A) or with anti-HBcAg antibody (B) followed by negative staining and observation in an electron microscope.

**Suppl. Fig. 5:**
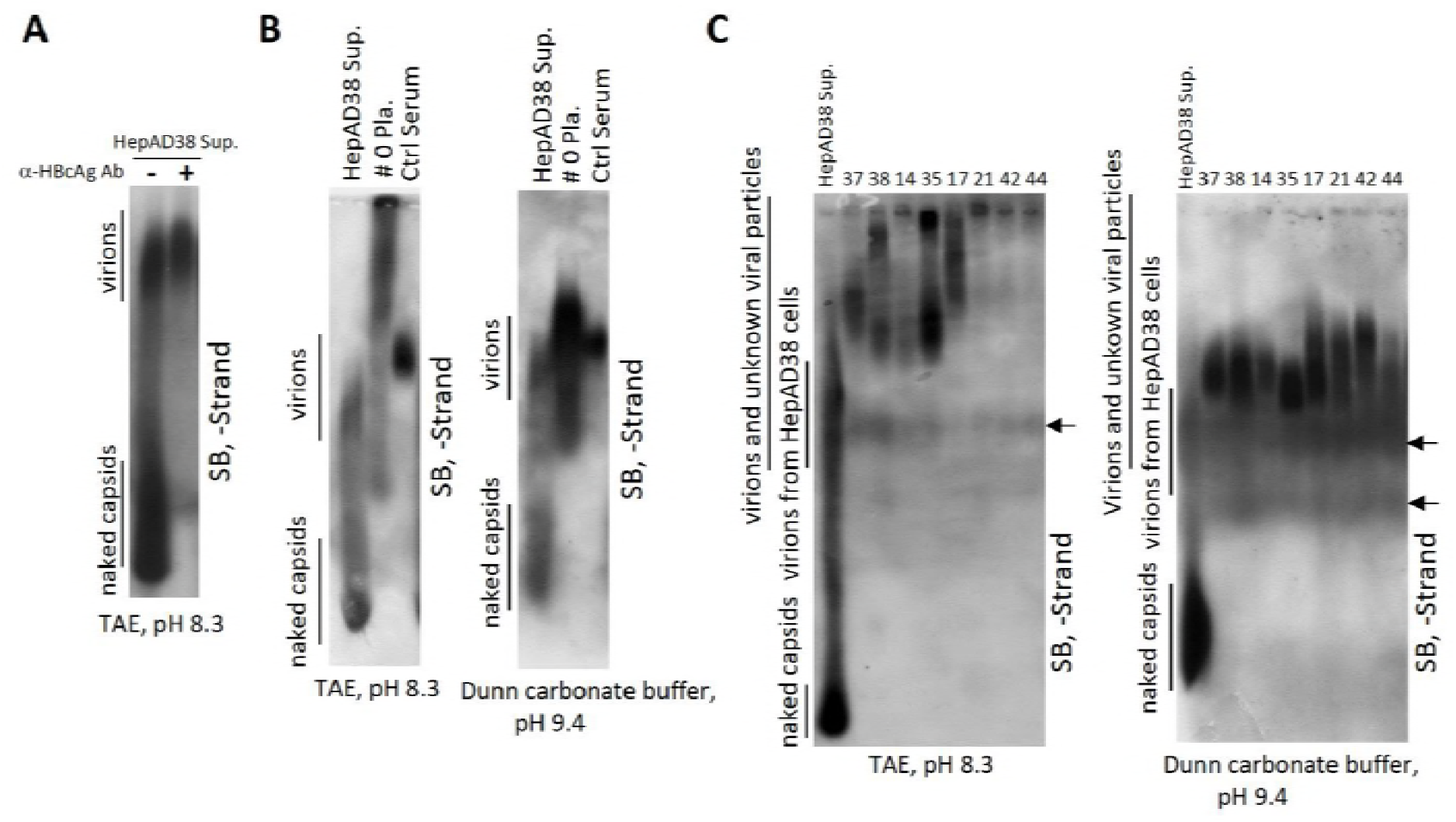
Native agarose gel analysis of viral particles in sera from hepatitis B patients. Ten microliter of HepAD38 cell culture supernatant (concentrated by ultrafiltration) incubated with or without anti-HBcAg antibody was resolved by native (TAE) agarose gel (0.8%) electrophoresis followed by Southern blot analysis with minus-strand specific riboprobe (A). Ten microliter of concentrated HepAD38 cell culture supernatant, plasma sample of patient 0 (not concentrated) and serum of a chronic hepatitis B carrier without liver inflammation (not concentrated) were loaded into agarose gels prepared in TAE buffer (pH 8.3) (B, left) or Dunn carbonate buffer (10 mM NaCHO_3_, 3 mM N_2_CO_3_, pH 9.4) (B, right) and separated overnight. Viral particle-associated DNA was detected by Southern blot analysis. Sera from patients 37, 38, 14, 35, 17, 21, 42 and 44 (10 μl each) were resolved by electrophoresis through 0.7% high strength agarose (type IV agarose used for pulsed-field gel electrophoresis) gels either prepared in TAE (C, left) or Dunn carbonate buffer (C, right) followed by Southern blot analysis. Arrows indicated trace amount of viral DNA from unknown viral particles.

**Suppl. Fig. 6:**
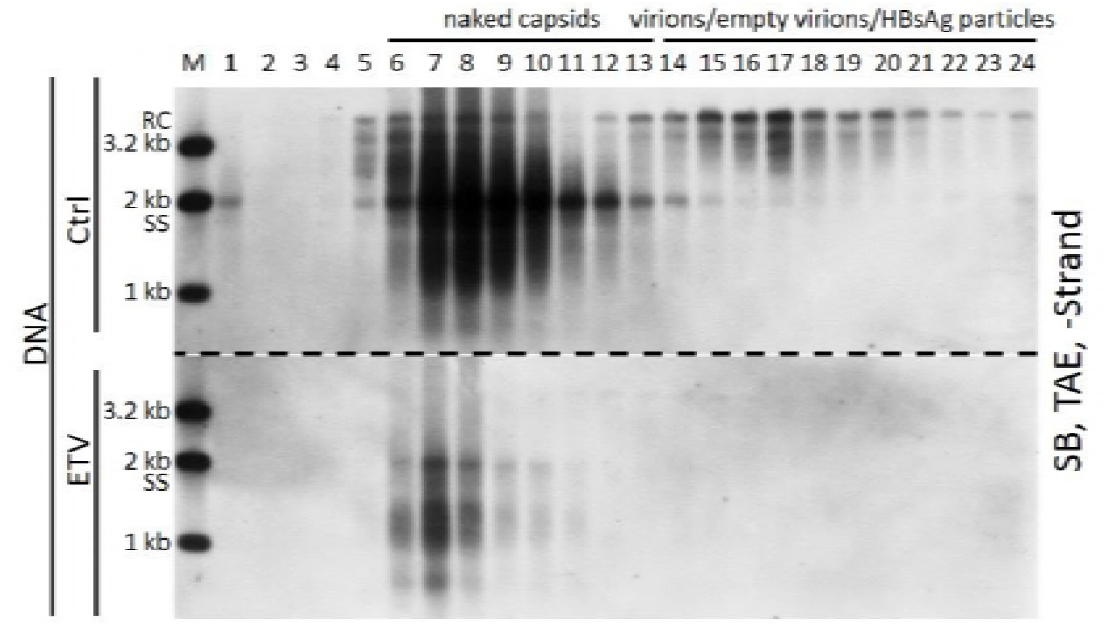
CsCl density gradient analysis of extracellular HBV DNA and RNA change upon entecavir treatment in HepAD38 cell culture supernatant. Longer exposure results of Fig 8G.

**Suppl. Fig. 7:**
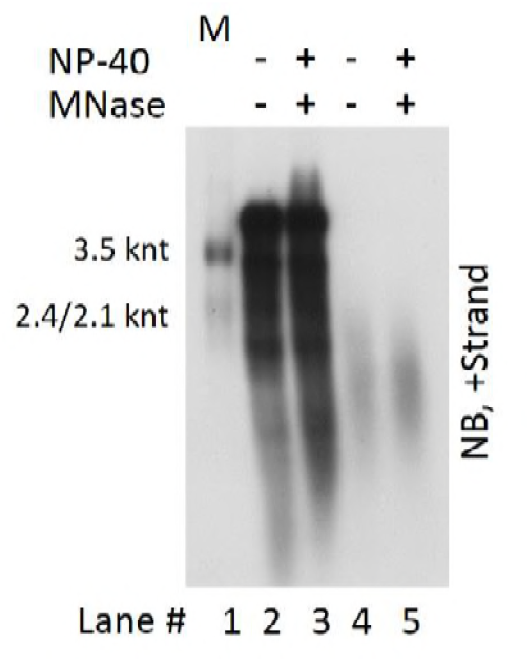
Change of extracellular HBV DNA and RNA upon detergent and micrococcal nuclease (MNase) treatment in HepAD38 cell culture supernatant. HepAD38 cell culture supernatant was untreated or treated with MNase in the presence of 1% NP-40. Reaction was stopped by addition of EDTA followed by digestion with proteinase K at 37°C for 30 min. Subsequently, viral DNA and RNA were released by addition of SDS (final concentration: 1%) and samples were incubated at 37°C for 15 min. Viral DNA and RNA mixture (lanes 2 and 3) or viral RNA (treated with DNase I [final concentration: 2 U/μl]) (lanes 4 and 5) were finally subjected to Northern blot analysis.

